# Z-Flipon Variants reveal the many roles of Z-DNA and Z-RNA in health and disease

**DOI:** 10.1101/2023.01.12.523822

**Authors:** Dmitry Umerenkov, Alan Herbert, Dmitrii Konovalov, Anna Danilova, Nazar Beknazarov, Vladimir Kokh, Aleksandr Fedorov, Maria Poptsova

## Abstract

Identifying roles for Z-flipons remains challenging given their dynamic nature. Here we perform genome-wide interrogation with the DNABERT transformer algorithm trained on experimentally identified Z-DNA sequences. We show Z-flipons are enriched in promoters and telomeres and overlap quantitative trait loci for RNA expression, RNA editing, splicing and disease associated variants. Surprisingly, many effects are mediated through Z-RNA formation. We describe Z-RNA motifs present in SCARF2, SMAD1 and CACNA1 transcripts and others in non-coding RNAs. We also provide evidence for another Z-RNA motif that likely enables an adaptive anti-viral intracellular defense through alternative splicing of KRAB domain zinc finger proteins. An analysis of OMIM and gnomAD predicted loss-of-function datasets reveals an overlap of predicted and experimentally validated Z-flipons with disease causing variants in 8.6% and 2.9% of mendelian disease genes respectively, with frameshift variants present in 22% of cases. The work greatly extends the number of phenotypes mapped to Z-flipon variants.

## Introduction

The discovery of the Zα domain in the p150 isoform of the double-stranded RNA (dsRNA) editing enzyme ADAR1 (encoded by ADAR), along with genetic studies in both humans ^1^ and mice ^2–4^ has unambiguously confirmed a biological role for both Z-DNA and Z-RNA (collectively called ZNA) in the regulation of interferon responses, self/nonself transcript discrimination ^5^ and the necroptosis cell death pathways ^6^. The covalent modifications of adenosine-to-inosine (A➔I) RNA editing performed by ADAR1 and the MLKL phosphorylation activated by ZBP1 (ZNA binding protein 1) enabled tracking of transient ZNA formation in cells. Here we use a genome-wide approach to discover additional phenotypes that are regulated by Z-flipons, sequences that can form ZNAs under physiological conditions. Our approach is computational and based on a novel and highly efficient algorithm for predicting Z-flipons based on experimental data. We leverage the large number of orthogonal datasets from the human genome and ENCODE projects to evaluate the validity of many hypotheses and present here those that are not falsified by existing experimental evidence.

We started with pretrained DNABERT model^7^ and fine-tuned it with validated Z-flipons from human genome-wide experimental studies (Figure 1). The resulting Z-DNABERT significantly outperformed previous approaches, such as DeepZ ^8^ that are based on convolutional and recurrent neural networks, with a recall of 0.89, precision of 0.78, and ROC AUC of 0.99 (Supplemental Table 1). The algorithm generates easily interpretable attention maps of Z-prone sequences at nucleotide resolution (Figure 1, Supplemental Figure 1).

**Figure 1.**
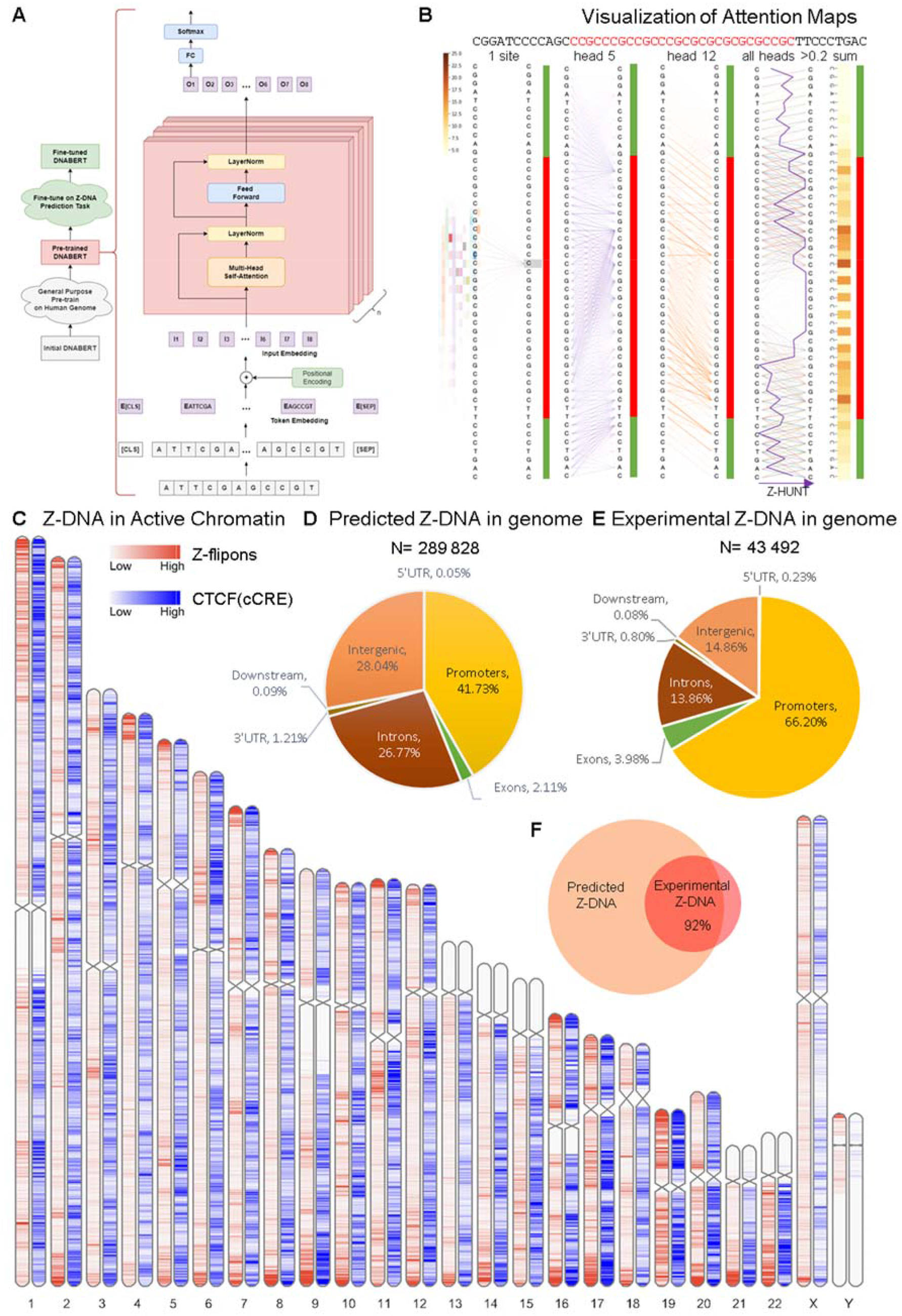
Generation of whole-genome Z-flipon maps with the Z-DNABERT model. **A.** Architecture of Z-DNABERT showing finetuning of DNABERT on experimental Z-DNA datasets. **B.** Interpretation of Z-DNABERT model. Visualization of attention scores for the sequence shown at the top of the panel that has the experimentally validated Z-DNA region colored red. From left to right: attention map for a single nucleotide; attention map from the head 5; attention map from head 12; attention map output that combines all layers with a threshold > 0.2;a line showing Z-hunt3 scores across the sequence; heatmap summarizing Z-DNA propensity. **C.** Whole-genome map with predicted Z-flipons compared to a map of CTCF protein binding sites that are present in candidate cis-regulatory elements (cCRE) defined by the ENCODE consortium **D.** Genomic features of the predicted Z-flipons. **E.** Genomic features of the experimental Z-flipons. **F.** Venn diagram of the overlap between predicted and experimental Z-flipons.

A large number of datasets are available for mapping DNA variants to phenotype, enabling us to perform a deep analysis of how flipons encode genetic information. We restricted this work to regions of experimentally verified Z-DNA, focusing on those overlapping genomic variants previously identified by Genome-wide Association Studies (GWAS) and disease focused approaches. We then performed computational mutagenesis with Z-DNABERT to test directly whether SNP alleles affected Z-DNA formation, then used haplotype analysis to map flipon alleles to trait values. We also assessed the role of Z-flipons in mendelian disease. Our findings expand the range of phenotypes attributable to Z-flipons beyond the human mendelian type I interferonopathies caused by loss of function (LOF) ADAR1 p150 variants ^1^.

## Results

### Developing generalizable deep learning model for Z-DNA prediction

Currently there are two human experimental datasets available that provide information on Z-DNA formation within human cells: the Shin et al ChIP-seq (Chromatin Immunoprecipitation followed by DNA sequencing of fragments) experiments with a resolution of 100-150 basepairs (bp) ^9^ and the nucleotide resolution, permanganate/S1 nuclease dataset (KEx) from Kouzine at al. ^10^.

For the deep learning model, we chose DNABERT pretrained with 6-mers representation. The approach is based on the Bidirectional Encoder Representations from Transformers (BERT) algorithm ^7^. We then trained the model further using the experimental datasets to create Z-DNABERT (Figure 1A, Methods and Supplemental Methods). We compared performance of Z-DNABERT with two other machine learning methods: DeepZ ^8^ and Gradient Boosting (CatBoost realization) ^11^. The latter approach also learns from k-mers representation (Supplemental Table 1). Z-DNABERT showed high performance on F1 and ROC AUC when tuned with the large nucleotide resolution KEx set. Part of the reason is shown by the Shin et al analysis where attention can is paid to poor ZNA forming sequences such as AAAAAA that are also enriched in the small number of 100-150 bp fragments analyzed (Supplemental Table 2). We used the KEx-tuned model for the work presented here.

Z-DNABERT outputs attention maps that are easily visualized (Figure 1B, Supplemental Figure 1). One can analyze output summarized for all heads or that for a particular head. Unlike the black box results from neural nets, the zebra-stripe patterns produced are easily interpretable: they show the propensity of alternating purine/pyrimidine dinucleotide repeats to form Z-DNA. The dark stripes correspond to purine bases that flip from the *anti* to the *syn* conformation as the transition from the right-handed to the left-handed helix occurs. The preference for guanosine over adenosine and cytosines over thymidine reflects the experimentally determined *in vitro* energetics that the -HUNT3 program uses to score Z-prone sequences ^12^. Compared to the Z-HUNT3 output (“all-heads” column Figure 1B), attention maps provide extra information on the sequence dependence of B-Z junctions rather than assigning them a fixed energy cost. These additional details likely account for the slight differences in predicted ranking of Z-prone motifs compared to the experimental Z-DNA input data (Supplemental Table 2). The Z-DNABERT model trained on human data also performed well in predicting Z-prone sequences from the mouse genome (Supplemental Table 3). Z-DNABERT also can predict the effect on Z-DNA formation of substituting any nucleotide in a sequence with another.

### Whole-genome prediction of Z-flipons

With Z-DNABERT trained on nucleotide resolution KEx, we generated genome-wide whole genome maps of Z-DNA regions (Figure 1C and Supplemental Data 1-4), which resulted in 290,071 regions covering 3167809 bp (0,16%) of the hg38 genome build. The genomic coverage of predicted Z-flipons was much more extensive than that for KEx (Figure 1C versus Supplemental Figure 2). We observed colocalization of Z-flipons with candidate cis regulatory elements (cCRE) defined by the ENCODE Consortium in many regions, with a higher density of overlaps in sub-telomeric regions. The correspondence with CTCF(CCCTC-binding factor) enriched sites at cCRE promoters is quite evident (Figure 1C) ^13^ and more pronounced than when each feature is considered separately (Supplemental Figure 3). Around 30% of the predicted Z-flipons fell within promoters and were less than 1 kb from a transcription start sites (TSS), with around 40% less than 3 kb distant. 30% are located in the introns with 7% found in the first introns and another 30% comprise intergenic regions. The enrichment of Z-flipons in promoter regions is consistent with previous analyses (Figure 1D) ^14^. Overall, the predicted Z-flipon set incorporates 92% of experimentally validated Z-DNA (Figure 1E and Supplemental Figure 2), but is 7 times larger. The maximum overlap of experimental Z-DNA vs predicted (95.32%) is observed in 5’ exons <300 bp from the TSS (Supplemental Table 4). We did not detect substantial overlap with regions of G-banding or with high recombination (Supplemental Figure 3) ^15^.

### Z-flipons are enriched in CTCF-bound proximal enhancer and promoter regions

We explored the cCRE results presented in Figure 1C further. Almost 10% of the predicted Z-DNA fell into cCRE regions (91,292 out of 926,535). Specifically, enrichment was observed in CTCF-bound proximal enhancer (3-fold enrichment) and promoter (6.7 fold enrichment) regions (Figure 2A, Supplemental Data 1), consistent with a regulatory role for Z-flipons.

**Figure 2.**
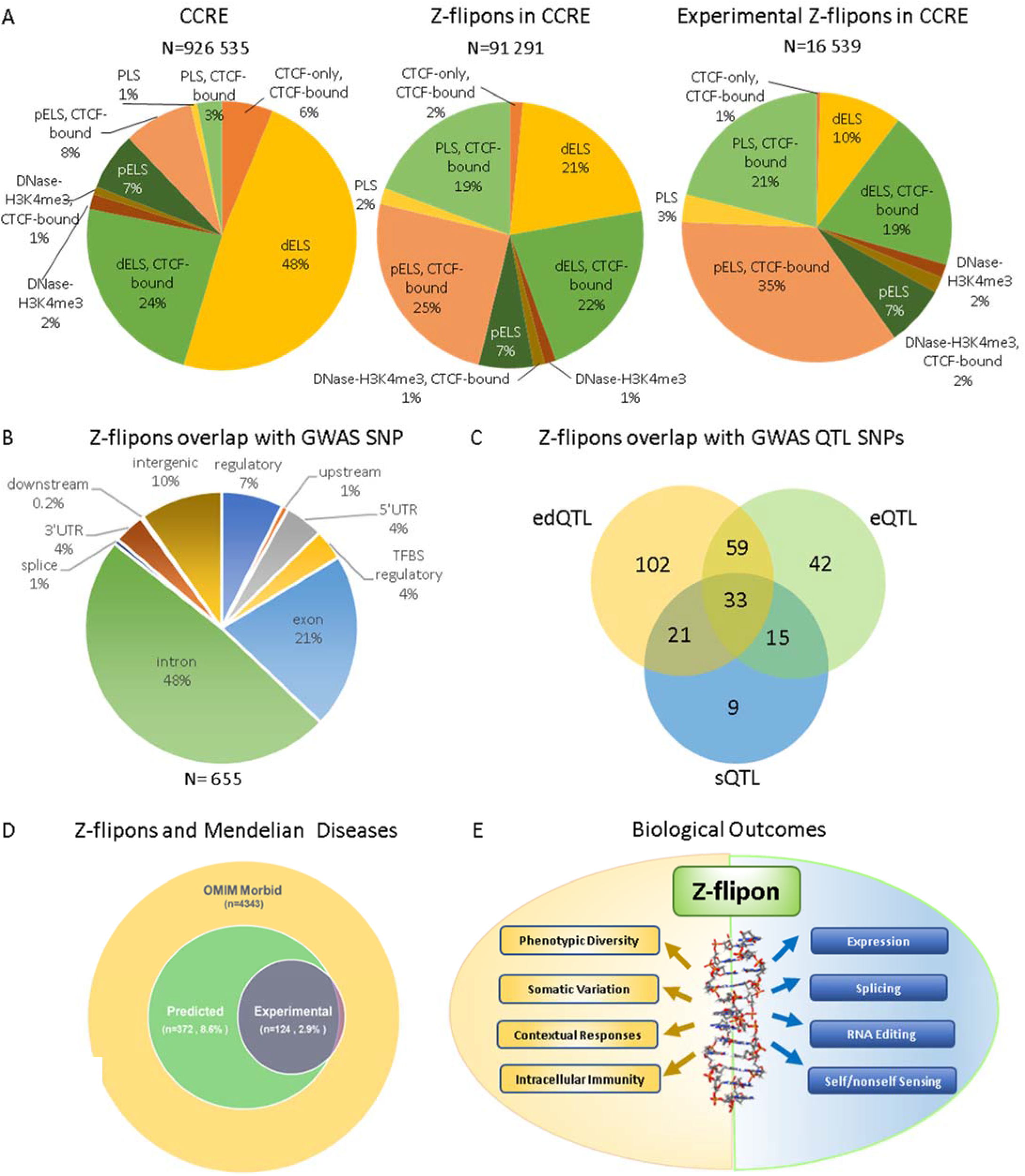
Z-flipon overlaps with orthogonal genomic data. **A.** Z-flipon overlaps with cCRE. **B.** Z-flipon overlaps with SNPs from the GWAS catalog. **C.** Predicted and experimental Z-flipon overlaps with GWAS QTL SNPs. **D.** Genes in the OMIM Morbid database with variants that overlap predicted and experimental Z-flipons. The predicted Z-flipon set caputures118 of the 124 genes with variants overlapping experimentally validated -flipons. **E.** The many ways that Z-flipons impact phenotype.

There were 393 of these transcription-associated cCRE regions where Z-flipons overlapped variants identified by GWAS. Among them, 86 (22%) are editing quantitative trait loci (edQTL) variants, 66 (17%) are expression QTL (eQTL) and 29 (7%) are splicing QTL (sQTL). Some of the QTLs are quite distant from the site of their effect, with some reported as more than 400 kb from an affected RNA editing site. Such a distance between associated elements suggests that Z-flipons can act by altering the loop topology of chromatin domains to bring widely separated elements close together, facilitating their interaction ^16,17^.

### Overlap of Quantitative Trait Loci with Z-flipons

We overlapped predicted Z-flipons with disease associated variants from the GWAS catalog (Figure 2B and Supplemental Data 2). We observed 3.2-fold enrichment of GWAS single nucleotide polymorphisms (SNP) in Z-flipons. Out of 108,517 unique GWAS SNPs. 655 (0.6%) fell into Z-DNA regions. We compared experimental Z-DNA predictions with respect to overlap with GWAS variants, and found that Z-DNABERT predicts 95% (109 out of 115) variants from KEx. Expanding the GWAS associated region by 500 or 1000 bases either side further increased the overlap with Z-DNABERT hits to 12440 and 20171 respectively (Supplemental Data 2).

We examined the overlap of Z-flipons with GWAS variants that are also QTLs for editing levels, expression level or splicing (Supplemental Data 2). Out of 661 variants from GWAS overlapping ZDNABERT, 215 (33 %) are edQTL, 149 variants (23%) are expression eQTL, and 78 variants (12%) are sQTL (Supplemental Data 2). We explored GO enrichment of variant falling in Z-flipons and found enrichment in positive regulation of transcription from RNA polymerase II promoter (GO:0045944 FDR = 2.78E^-04^) and chromatin (GO:0000785 FDR = 7.89 E^-03^) consistent with our other findings.

There was also a significant overlap of Z-flipons in the OMIM collection of mendelian variants (Figure 2D) that we will discuss later as we develop the evidence for the flipon dependent outcomes summarized in Figure 2E.

### Z-flipons in Action

A natural question is to ask how flipon variants affect trait values. Our analyses identify two novel repeat motifs involved in expression and splicing, both differing from the conserved Alu Z-Box motif we previously identified as targeting A➔I editing by the ADAR1 p150 isoform ^18^. The first motif has a Z-RNA stem associated with loop containing an effector domain and the other represents a previously characterized intronic splicing enhancer sequence that can also fold into a Z-RNA helix.

#### An eQTL in SCARF2 affects MED15 and Height

The rs874100 SNP (NM_153334.7:c.2459G>C), which encodes a nonsynonymous variant (NP_699165.3:p.Gly820Ala) (Figure 3A), overlaps a predicted and experimentally confirmed Z-flipon (Figure 3B). The Z-DNABERT mutagenesis map reveals that the minor C allele disrupts Z-DNA formation (Figure 3C). The allele also prevents the fold of the SCARF2 transcript into Z-RNA (Figure 3D). The fold forms a loop anchored by the Z-RNA stem that contains a GU splice donor site at position 90, although there is no current evidence that the site is associated with alternative splicing.

**Figure 3.**
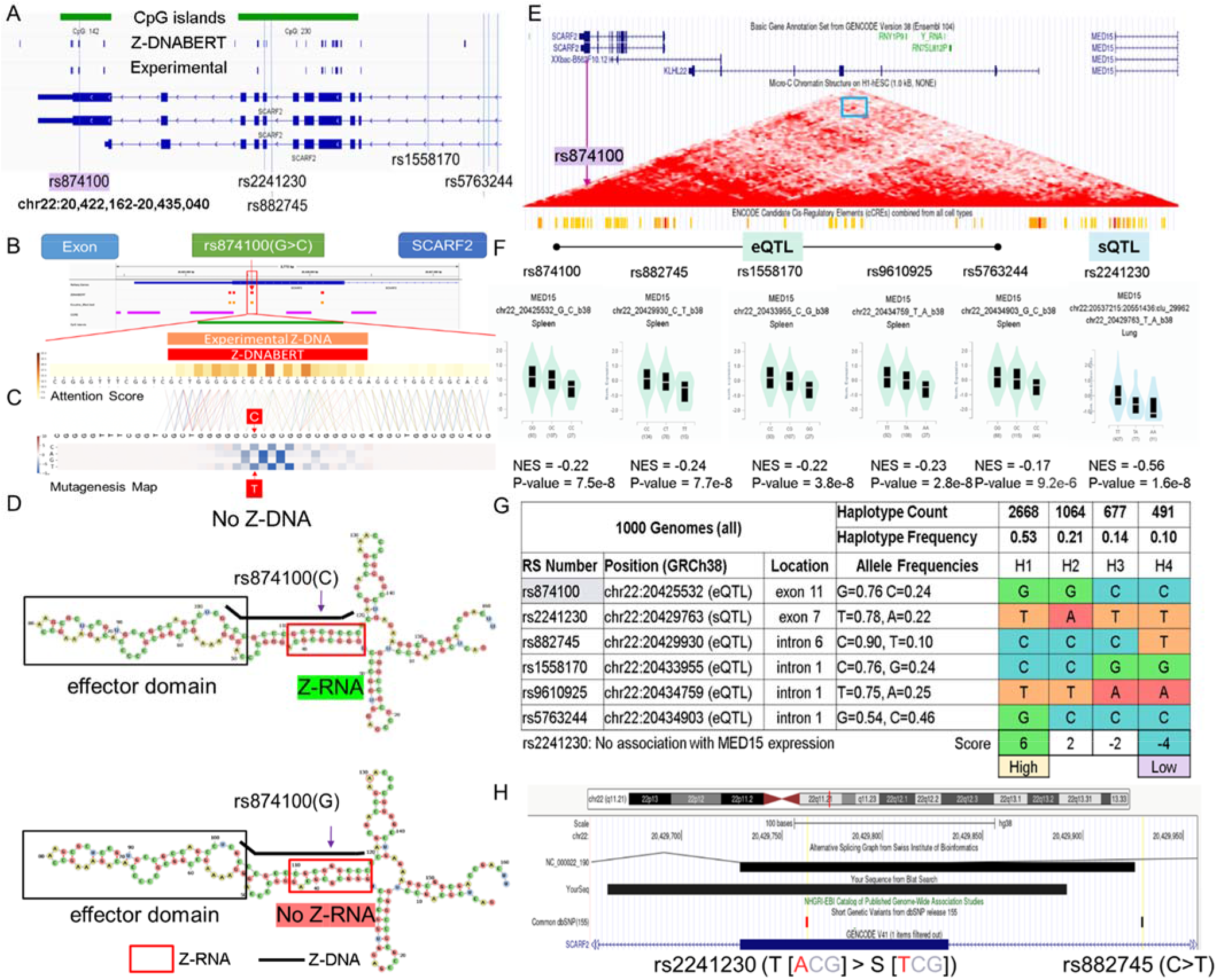
A Z-flipon in SCARF2 associates with MED15 expression **A.** chr22:20,422,162-20,435,040 showing the 3’ region of SCARF2 along with SNPs used in the analysis. The position of predicted and experimentally confirmed Z-flipons are also shown, along with CpG islands. **B.** Overlap of eQTL rs874100 with Z-DNABERT Z-DNA prediction **C.** Computation prediction of the effect of mutagenesis of each nucleotide in the Z-flipon region. The SNP variant A allele leads to loss of Z-DNA formation**. D.** The Z-RNA fold with the Z-DNABERT Z-DNA sequence is shown below the thick black line. Note that the RNA is transcribed in the reverse direction from the genome. The SNP minor allele also disrupts the Z-RNA fold. **E**. chr22:20,415,440-20,518,466 showing both SCARF2 and MED15 genes, along with a microC map from human embryonic stem cells (HESC), with the blue box highlighting the convergence of the red diagonals that indicate contacts between rs874100 and the MED15 promoter. The orange bars show the cCRE in the MED15 promoter that were mapped by the ENCODE consortium. **F.** SNP eQTL for MED15 showing the normalized effect size (NES) and p-value determined by the GTEX consortium. **G.** The haplotypes were scored by adding +1 if a SNP allele was associated with an increase in trait value and −1 if the value was lower. Haplotype 1 favors Z-DNA formation and is associated with high MED15 expression while Haplotype 4 has low expression of MED15 and a low propensity to form ZNAs. **H**. The rs2241230 SNP is positioned near an alternative splice site forSCARF2 and is a sQTL for MED15.

The rs874100 SNP is an eQTL for the mediator complex subunit 15 (MED15) gene that is associated by GWAS with height. The microC map from human embryonic stem cells (hESC) reveals the presence of contacts between the rs874100 region and the MED15 promoter (Blue Box, Figure 3E). We were able to define four haplotypes that incorporate other neighborhood SNPs that are also associated with height (Figure 3F, G). The haplotypes also included the exon 7 nonsynonymous SNP rs2241230 (NM_153334.7:c.1273A>T variant (XP_016884554.1:p.Thr425Ser), which is not an eQTL but rather a sQTL and the intron 6 variant rs882745 (NM_153334.7:c.1203-97G>T) that is just upstream of an alternative splice site for SCARF2 (Figure 3H). We scored the haplotypes and identify those associated with high and low expression of MED15.

The ZNA prone haplotype H1 is associated with increased expression of MED15 while haplotype H4 with the rs874100 C allele that disrupts the Z-DNA stem has low expression. This finding is supported by the two intronic SNPs rs1558170 (NC_000022.11:g.20433955C>G) and rs9610925 (NC_000022.10:g.20789046T>A) that are in strong linkage disequilibrium with rs874100. The increased MED15 gene expression of H1 relative to H4 could partly reflect the nonsynonymous changes produced by the SCARF2 SNPs rather than through differences in Z-DNA formation. This explanation is less likely as the rs874100 amino acid substitution has been shown by clinical testing to be benign (ClinVar accession RCV000602615.1). Further, the variant produced lies in the disordered carboxy terminus of the protein and not within a functional domain. The other nonsynonymous SNP rs2241230, is not an eQTL for MED15, but a sQTL whose minor allele is associated with decreased splicing of MED15, likely offsetting the increased expression associated with the rs874100 G allele. The association of rs874100 with height may then reflect the higher expression of MED15 protein due to the formation of ZNAs by H1. The increased coupling between enhancers and promoters would increase cell growth by generating higher levels of transcripts and proteins. The altered splicing associated with rs2241230 may further affect MED15 expression levels by altering the isoforms produced.

#### An eQTL in SMAD1 affects HDL cholesterol

We observed a similar Z-RNA stem/loop motif in our analysis of eQTLs for SMAD1. The eQTLs present in the 5’UTR of the SMAD1 gene include rs13144151(A>G) (NC_000004.11:g.146403165A>G) and rs13118865(C>T) (NG_042284.1:g.5698C>T). The SNPs defined three haplotypes that express intermediate (H1), high (H2), and low (H3) levels of SMAD1 mRNA. Both H2 and H3 contain potential Z-RNA forming sequences. The high expressing H2 incorporates the minor G allele of rs13144151 that overlaps an experimentally validated Z-DNABERT prediction (Figure 4D). Mutagenesis mapping of rs13144151 with Z-DNABERT revealed that G allele caused a slight increase in Z-propensity. While not pronounced at the level of DNA, the effects of the allele on the RNA fold are quite evident (Figure 4G, H): the G allele stabilizes an additional potential Z-RNA helix by adding an extra G:C bp to increase its span to 6 bps, producing the minimal length substrate required to dock a Zα domain ^19^ (Figure 4G, H). The low expressing H3 haplotype is defined by the minor alleles of rs13118865 and rs1264670 (G>A) (NC_000004.11:g.146402927G>A). Rs1264670 is incorporated into an RNA fold motif similar to that of H1 with a Z-RNA stem and a hairpin loop domain (Figure 4I). Present in the domain are two unpaired splice donor sites and many CGGG sites of the type bound by the alternative splicing factor RBM4 (RNA binding motif protein 4). Since RBM4 is known to suppress use of splice donor sites ^20^, we refer to the hairpin as an effector domain.

**Figure 4.**
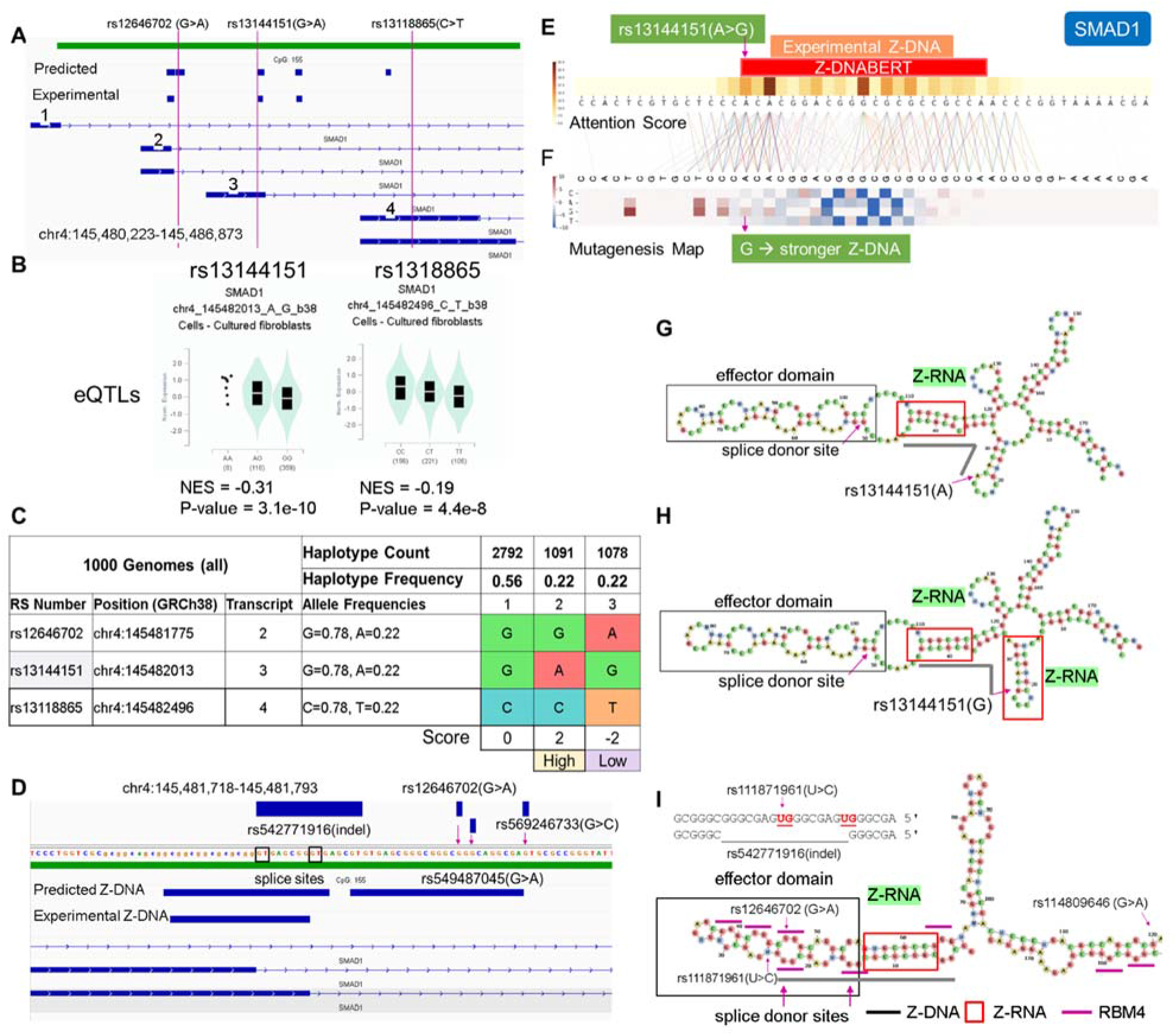
SMAD1 expression and splicing for rs13144151(A>G) and rs12646702(G>A) **A**. Location of SNPs and splicing isoforms. The exons that are labeled 2, 3 and 4 are associated with different transcripts. Ater splicing, each transcript is uniquely marked by the presence or absence of a particular SNP in one of the numbered exons. **B** The rs13144151(A>G) and the rs13118865(C>T) SNPs affect expression of SMAD1 mRNA. No QTL data is available for rs12646702, but it is in linkage disequilibrium with rs13118865 that serves as a surrogate. **C.** Haplotypes differ in their expression of SMAD1. The haplotypes were scored by assigning +1 to the alleles that increased trait values and −1 otherwise. For rs12646702 where no quantitative trait information is available, both alleles were assigned a value of zero. **D.** The 5’ UTR of SMAD1 in the vicinity of rs1264670(G>A) showing the Z-DNABERT predicted Z-flipons, experimental Z-flipons, SNPs and an alternatively spliced SMAD1 exon. **E**. Mapping at nucleotide resolution of the overlap of Z-DNABERT predictions and rs13144151 **F.** Z-DNABERT predicted effects of nucleotide substitutions at this locus showing that the A>G substitution enhances Z-DNA formation. **G**. The Z-RNA stem and the effector domain loop containing a splice donor site formed in the vicinity of rs1314415. The heavy black line corresponds to the Z-flipon sequence predicted by Z-DNABERT. **H.** The rs1314415 G allele enables formation of an additional Z-RNA stem that is associated with lower expression of the transcript. **I**. Z-RNA forming stem that includes rs12646702 is associated with an effector domain that contains CGGG binding sites for the alternative splicing factor RBM4 indicated by short purple lines, with the heavy black showing the Z-flipon sequence. The SNP locations are shown along with the rs542771916 indel.

Interestingly, the SNP minor alleles affecting SMAD1 expression map not only to haplotypes, but also to the exons defining different splice isoforms. The rs13144151 A allele that defines the H2 haplotype is present on exon 3, while H3 is defined by both the rs13118865 T allele on exon 4 and the rs1264670 A allele at the 3’ end of exon 2 (as labeled in Figure 4A). The strength of Z-RNA formation associated with each exon likely affects the expression of each isoform. The transcription of isoforms containing exon 3 may be favored by the rs13144151 A allele that disrupts Z-RNA formation and allows RNA polymerase progression. In contrast, both exons 2 and 4 contain strong Z-RNA folds that could cause RNA polymerases, leading to lower readout of these isoforms.

The association of rs13144151 with HDL cholesterol levels (A allele = −0.018 unit decrease ^21^) is consistent with the known role of SMAD1 in negatively regulating cholesterol efflux from cells. Increased SMAD1 expression leads to decreased levels of the cholesterol transporters ABCA1 and ABCG1, with lipid accumulation by macrophages producing foam cells that are associated with atherosclerosis ^22^.

### A sQTL in CACNA1C affects DCP1B and BMI

We also assessed the relative roles of Z-DNA and Z-RNA in splicing by analyzing sQTLs found in the 5’ UTR of CACNA1C (calcium voltage-gated channel subunit alpha1 C) that alter processing of decapping1B protein (DCP1B) transcripts, DCP1B protein initiates mRNA decay by enzymatically removing the 5’ cap from RNAs. The index SNP rs11062091 (NG_008801.2:g.87418G>A) is a sQTL for DCP1B splicing but is not currently with a phenotype. Of the SNPS in the region nearby, rs2470397 (NG_008801.2:g.31165T>C) is a sQTL associated with BMI In a GWAS of BMI in nearly half a million individuals ^23^; rs10774018 (NG_008801.2:g.82974G>C) is a sQTL associated with visceral obesity and height ^24,25^ and rs2108635 (NG_008801.2:g.84605A>G) is associated with BMI but is not a QTL (Figure 5A). Haplotype analysis revealed that the major allele of rs11062091 is on a haplotype H3, which scored highest for splicing, while the minor allele is on H6, which has the lowest score. In these haplotypes, rs2470397 alleles are not correlated with those of other SNPs, reflecting the high recombination rate recorded for this chromosomal segment (Figure 5B). Nevertheless, the rs2470397 minor C allele helps define haplotypes 3 and 6 and the association of the rs11062091 minor A allele with low DCP1B splicing and increased obesity (Figure 5B). The effect on BMI may reflect the rate at which transcripts undergo decay, with H6 increasing the longevity of transcripts that promote adiposity.

**Figure 5.**
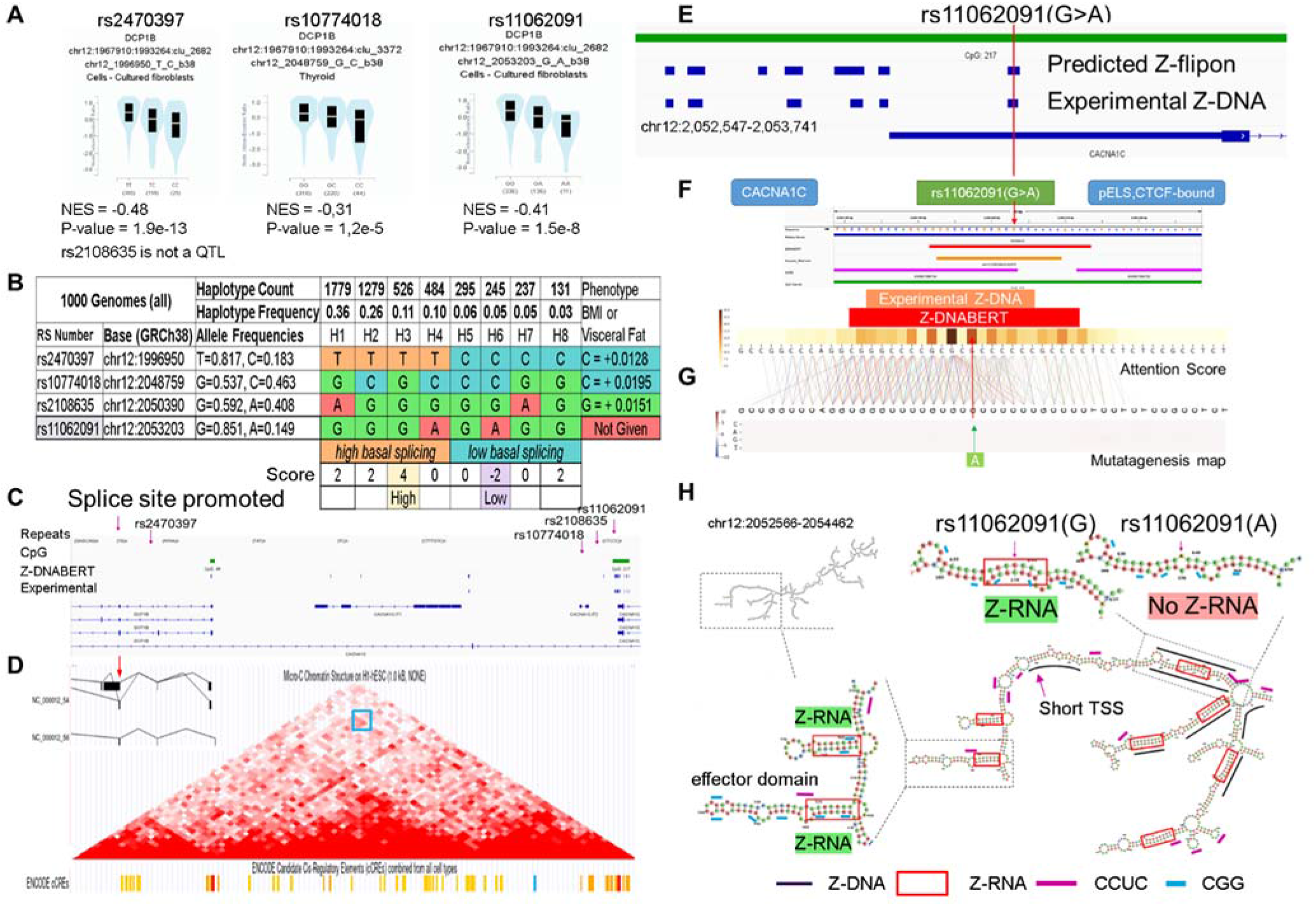
Z-flipons in (hg38.chr12:1,935,235-2,714,656) in CACNA1 (calcium voltage-gated channel subunit alpha1 C) affect splicing of DCP1B (decapping mRNA 1B) **A.** Minor SNP alleles are associated with decreased splicing of DCP1B transcripts **B.** Haplotype map of the region that supports an association between decreased DCP1B splicing and increased body mass index. The haplotypes were scored by adding +1 to the total if the allele was associated with an increase in trait value and −1 if the value was lower. The highest and lowest scores are associated with rs11062091 alleles. **C.** Location of the alternative DCP1B splice along with the position of all SNPs. **D.** The alternatively spliced DCP1B transcript is drawn as an inset to the microC map that shows contact is present between the SNP locus and the DCP1B genic region, as indicated by the region boxed in blue. The areas of contact contain chromatin modifications classified as cCRE by the ENCODE project (orange bars represent enhancers and red bars are for promoters). SNP positions, simple repeats and both predicted and experimental Z-DNA are shown. The CACNA1C splice site affected by rs11062091 is upstream (chr12:1967910-1993264) and is currently not annotated in GENCODE 41. **E.** Expanded view of Z-DNA in the vicinity of rs11062091 showing the overlap between the Z-DNABERT predicted and experimentally validated Z-flipon **F.** Z-DNABERT prediction for the Z-flipon that incorporates rs11062091. **G.** -DNABERT mutagenesis shows that single nucleotide variants do not affect the propensity of the rs11062091 Z-flipon to form Z-DNA. **H.** Progressively zoomed in views of the dsRNA fold of the transcript from the rs11062091 region. The A allele of rs11062091 disrupts formation of Z-RNA. The black lines show the experimental determined regions of Z-DNA formation. Only the rs11062091 Z-flipon experimentally forms Z-DNA at the locations where the two RNA strands that create the Z-RNA stem are transcribed from. Multiple Z-RNA prone helices are formed with RNAs transcribed from regions where Z-DNA formation was not experimentally detected. The short purple lines show CCUC motifs that could represent CTCF protein binding sites. The RNA fold overlaps the transcription start site (TSS, chr12:2,052,986) for the shorter CACNA1C transcript as indicated by the TSS label. A Z-RNA stem/loop effector domain motif resembling those in Figures 3 and 4 is also illustrated with short blue dashes above CGG repeat sequences.

The microC map from hESC showed contact between the region containing the alternative DCP1B splice site and rs11062091, both of which bear enhancer CCRE marks and an overlap with CTCF binding sites (Figure 5C and D). The region around rs11062091 has many predicted and experimental Z-flipons, yet Z-DNABERT mutagenesis maps revealed little effect of the SNP alleles on Z-DNA formation (Figure 5E). Analysis of the RNA fold revealed many regions of likely Z-RNA formation (red boxes) that did not align with experimentally validated Z-DNA (identified by heavy black lines). One of these contain a Z-RNA stem loop motif similar to those observed with SCARF2 and SMAD1 (Figure 5H).

With other Z-RNA stems, experimentally validated Z-DNAs aligned only with the upstream strand of the RNA and not its downstream complement. The only region where Z-DNA overlapped with both Z-RNA strands was the one that included rs11062091. The effect of the rs11062091 minor A allele was to disrupt formation of this particular Z-RNA helical stem (Figure 5H). The results suggest that the two Z-DNA elements producing the rs11062091 Z-RNA nucleate the remaining RNA fold. They then provide an anchor to promote seed a spliceosome condensate. Indeed, rs11062091 is a sQTL for RP5-1096D14.6 and CACNA1C-IT2 in addition to DCP1B. The 12 canonical CCTC motifs in Z-RNA associated effector domains could actively promote spliceosome formation by localizing CTCF to the region(Figure 5H) ^26^. Similar interactions may contribute shown in Figure 1. the alignment of CTCF/cCRE regions.

The failure of Z-DNABERT to detect many of the Z-RNA in this fold may reflect that the experimental determination of Z-DNA was performed in a single cell line. Alternatively, the result may be due to the different energetics of Z-RNA formation compared with Z-DNA. The preformed RNA bulges and bp mismatches in dsRNA facilitate A-Z creation ^27^, with each costing less energy than the 5 kcal/mol required for each B-Z-DNA junction. In the case of Z-RNA, formation of only a 6 bp binding site is required to dock the Zα domain ^19^. This length is much shorter length than Z-DNABERT is trained to discover (Supplemental Figure 2) as we require at least 11 contiguous bp.

### Z-flipons, Edited and Noncoding Transcripts

The Z-DNABERT training limited our exploration of the local effects of Z-flipons within genes that contain known sites of A➔I RNA editing. Indeed, Z-DNABERT does not detect the Z-RNA forming ALU sequencing known to impact ADAR1 editing of the cathepsin S (CTSS) RNA ^27,28^. We also did not expect the KEx dataset that analyzed Z-DNA to identify folds relevant to Z-RNA editing.

Overall, we found very few cases of direct overlap of Z-flipons with editing sites in the analysis of a number of published datasets (Supplemental Data 3). As many editing substrates are long, we also searched for Z-flipons in the 1kb surrounding the editing site and found a higher overlap. One experimental study explored editing suberates recognized by the Zα domain of ADAR p150 ^29^. Of the 1248 mRNAs identified, none had a Z-DNABERT overlap. Expanding the search window for a Z-flipon prediction to 1kb revealed that only 4% of the ADAR1 p150 editing sites overlapped. A separate study of lung adenocarcinoma tumors ^30^ found 1413 genes where the total level of RNA editing and expression were correlated. Of these, 5% of edited sites have a direct overlap with Z-flipons (Supplemental Data 3). Expanding the region of search for Z-flipons within 1kb of editing sites yielded a 19% overlap. We further found that 182 of the transcripts immunoprecipitated with the ZNA specific antibody Z22 from mouse embryonic fibroblasts ^6^ (Supplemental Data 3), providing some experimental evidence suggestive of Z-flipon conservation between mouse and human. For the ADeditome database, which maps 1,676,363 editing sites associated with Alzheimer’s Disease, only 271 overlapped a Z-flipon prediction, of which 6 were validated experimentally (Supplemental Table 5). In contrast Z-flipons were found within 1kb of editing sites in 50% of ADeditome genes (Supplemental Table 6).

The cases where we were able to overlap Z-flipons with editing sites were for Z-RNA stems 12 bp or longer (Supplemental Figures 6,7). The STXBP5L intronic dsRNA identifies in that manner was short and heavily edited (Figure 6). In contrast only a single edit (reproduced in the lung adenocarcinoma dataset(Supplemental Data 3 and in the Rediportal database) is present in the BIRCA transcript, raising the question of whether the edit is functional or whether it indicates that binding ADAR1 p150 has other outcomes. Interestingly this site is targeted by hsa-miR-8485 (and potentially by hsa-miR-574-5p and hsa-miR-297) that is bound by TDP-43 (encoded by TARDBP) to regulate a number of outcomes ^31^ raising the possibility that ADAR1 p150 regulates the access of hsa-miR-8485 to the BICRA transcript. Another instance where Z-RNA may enable regulation of noncoding RNAs is provided by RMRP (RNA component of mitochondrial RNA processing endoribonuclease) (Supplemental Figure 8) that performs many different functions through interactions involving miRNAs ^32^

**Figure 6:**
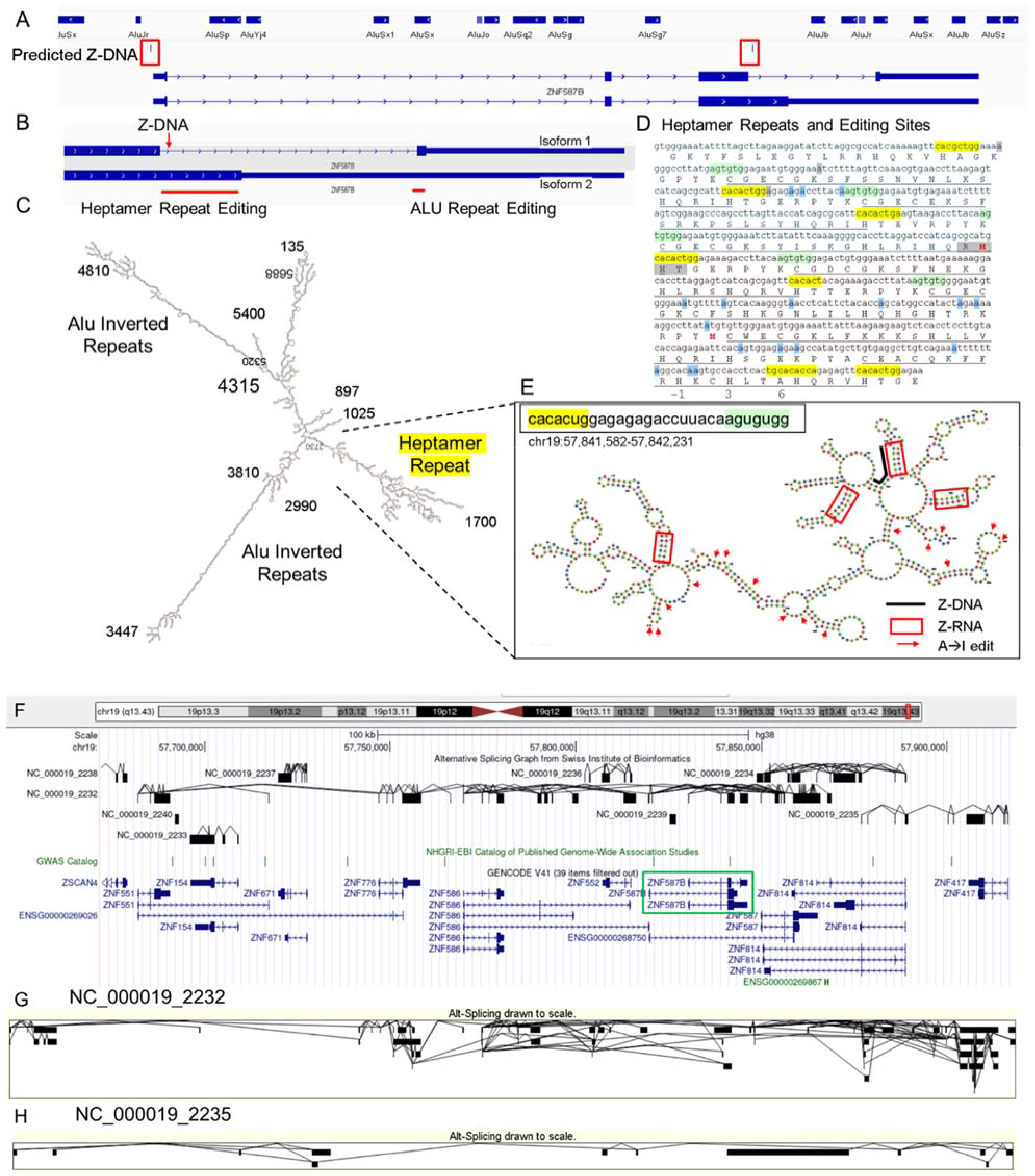
Nonsynonymous RNA editing of ZNF587B. **A** ZNF587B locus **B** ZNF597 isoform-specific RNA edits occur in different exons. **C.** dsRNA fold showing two classes of editing substrate **D.** dsRNA region maps to C2H2 Zinc Finger (ZNF) repeats that have a CX_2-4_CX_12_HX_2_-_6_H motif (X is any amino acid) and are underlined. The ZNF domains are joined by a seven amino acid linker that is within the heptad repeat. The gray box lies underneath the Z-DNABERT predicted Z-DNA sequence and the blue boxes highlight residues with nonsynonymous edits. The numbering immediately below the sequence in panel D corresponds to the DNA binding residues of the α-helix of the ZNF above. **E.** Heptad repeat folds are highlighted and the Z-RNA prone sequences are within the red boxes. The arrows indicate A➔I editing sites. The heavy black line is above the predicted and experimentally validated Z-flipon sequence. **F**. Alternative splicing within chromosome 19 telomeric zinc finger gene cluster (hg38. chr19:57,672,145-57,921,020) with two of the trans-splicing isoforms displayed in **G** and **H.**

#### ZNF587B and RNA editing

A predicted and experimentally validated Z-flipon within ZNF587B gene that is associated with many nonsynonymous edits lead us to investigate the locus further (Figure 6). The gene is in one of the zinc finger (ZNF) gene clusters enriched on chromosome 19 (Supplemental Figure 9). Depending on how it is spliced, ZNF587B contains up to 13 zinc finger domains (ZNF), plus a KRAB (Krüppel associated box) domain of the kind thought to mediate repression of transposon repeat elements ^33^. ZNF587B RNA editing is promoted by a number of ALU inverted repeats (AIR) similar to those of known ADAR1 substrates (Figure 6A). They overlap the terminal exon of one RNA isoform and result in RNA recoding specific to that transcript (Figure 6B and C). A different type of RNA fold directs editing of the other ZNF587B splice isoform (Figure 6B). Interestingly, the dsRNA in this region forms from heptamer repeats (HR) that create clusters of unpaired RNA loops distinct from the long, linear AIR substrates (Figure 6C, D and E). The HR has purine-pyrimidine inverted repeats capable of forming short Z-prone dsRNA helices ^19,27^ that resemble those clusters we recently identified in mouse by immunoprecipitation with ZNA specific Z22 antibody ^6^.

The length of the HR is conserved. It encodes the linker between adjacent ZNF (Figure 6D). Interestingly, the CACA motif overlaps that of known intronic splicing enhancers ^34^, raising the possibility that Z-RNA formation by the HR modulates alternative splicing. The arrangement of ZNF in clusters may enable intergenic splicing to generate new combinations of ZNFs at the RNA level. Evidence for the alternative splicing and trans-splicing from the Swiss Institute of Bioinformatics curated dataset is shown in Figures 6F-H.

The generation of these novel transcripts would be favored by the interferon induction of the known Z-RNA binding proteins ADAR1 p150 and ZBP1. The non-synonymous edits scattered through the fold are consistent with Z-RNA dependent localization of p150 to these transcripts. None of the edits alter the three residues (called −1, 3 and 6 as numbered on the bottom line of Figure 6D) that are involved in DNA recognition by ZNF^35^, so do not change the specificity of the ZNF. The altered splicing rather than RNA editing may be the major outcome produced by ADAR1 p150 as binding of p150 to the Z-RNA helix would occlude the site and make it unavailable to the splicing machinery. Alternatively, the interaction could help direct the locus to a spliceosome condensate. The novel combinations of ZNF produced by alternative splicing could prevent the escape of recently recombined transposons and viruses from KRAB mediated suppression.

### Z-flipons, Mendelian Disease and LOF Variants

We also examined Z-flipons for association with mendelian disease (Supplemental Data 4 and Supplemental Figures 10-18) given the previous emphasis placed on Z-DNA as a cause of genomic instability ^36^. There is overlap between experimentally determined Z-flipons and mendelian variants in a number of genes including HBA1 (hemoglobinopathies), CDKN2A (Melanoma Susceptibility), MC1R (red hair color, melanoma), WNT1 (Osteogenesis Imperfecta, type xv), NPHS1 (Nephrotic Syndrome, Type 1), SOX10 (Waardenburg Syndrome, Type 2e), IDUA (Hurler-Scheie Syndrome), LAMB3 (Heterotaxy), IL17RC (Familial Candidiasis) and FOXL2 (Blepharophimosis, Ptosis, And Epicanthus Inversus, Type I), providing direct evidence that Z-flipons do influence trait variation. Predicted Z-flipons also overlap with a more extensive range of OMIM phenotypes. Examples include TERC, the telomerase RNA, TERT, TP53, LMNA, NKX2.5, HBA2 and NROB1. Overall, we found an overlap of mendelian disease-causing variants with predicted (n=372) and experimentally validated (n=124) Z-flipons in 8.6% and 2.9% of OMIM genes(n=4343) respectively (Figure 2D). The majority of events (71%) with experimentally validated Z-flipons were due to nonsynonymous variants that altered arginine codons in 22% of cases (Supplemental Figure 19) while 22% of variants were LOF frameshifts (Supplemental Figure 20). We also analyzed the 430,056 predicted LOF (pLOF) variants listed in the Genome Aggregation Database (gnomAD) that are distributed over 18749 unique genes ^37^. Of these, 4362 variants fell into predicted Z-flipons. Interestingly, of the 1160 variants present in the KEx dataset, 1093 (94.2%) are in the gnomAD-pLOF set. Frameshift deletions were also more frequent with Z-flipon overlaps compared to other Z-flipon LOF classes and compared to the entire gnomAD-pLOF variant collection (Supplemental Figure 21 and Supplemental Data 5). Overall, 637 of the 2614 Z-flipon LOF genes (24.7%) overlapping the gnomAD-pLOF have OMIM morbid phenotypes (n=4343), compared to 21.5% of the gnomAD-pLOF genes. Interestingly, the overlap of the Z-flipons present in the gnomAD-pLOF with OMIM genes is much higher than the actual number of Z-flipons recorded in OMIM. There is a 14.7% overlap of genes with gnomAD-pLOF predicted Z-flipon variants and a 3.9% overlap with genes containing experimentally validated Z-flipons (Supplemental Figure 22, Supplemental Data 5). GO analysis of Z-flipon mendelian variants annotated in OMIM showed enrichment for transcriptional activity, homeobox proteins and transforming growth factor regulators of the extra-cellular matrix (Supplemental Data 4).

## Discussion

Discovering the functional roles of Z-flipons and mapping the associated phenotypes is a challenging task, as previously noted ^38^. We used genome-wide data and computational experiments to genetically map flipons to QTLs and disease outcomes. We used a machine learning approach called Z-DNABERT to detect Z-flipons by tuning the transformer algorithm implemented in DNABERT ^7^ with experimentally validated Z-DNA forming sequences obtained from the human genome at nucleotide resolution. Z-DNABERT outperformed previous approaches based on neural networks and enabled the findings reported here. Z-DNABERT was also helpful in finding Z-RNAs but did not directly detect those Z-RNA that we and others have demonstrated experimentally where the dsRNA helix length is shorter than 12 nucleotides long ^6,18,27^. The difficulty derives from the different energetic requirements for Z-DNA formation compared to Z-RNA, especially in the cost of establishing the junction between left and right-handed helices. The penalty is lower for Z-RNA than for Z-DNA as loops and mismatches facilitate RNA junction formation. Also, as with any dsRNA, Z-RNA requires close proximity of two complementary sequences, something Z-DNABERT is not trained to find. Despite these limitations we were able to use the ability of Z-DNABERT to perform computational mutagenesis to distinguish between Z-DNA and Z-RNA dependent events. Overall, we associate experimentally validated Z-flipons with active promoters that we then link to quantitative and disease phenotypes through the analysis of orthogonal genome-wide datasets. The work furthers our understanding of flipon biology and establishes a community resource. The hypotheses generated are data-driven and open new lines of enquiry into the germline and somatic mechanisms that lead to QTL variation and disease. They provide additional insights into pathways that produce intracellular immunity against retroelements and pathogens. They also suggest a role for Z-RNAs in regulating the interactions of noncoding RNA with other transcripts (Figure 2E) and establish a close connection between Z-flipons, CTCF and loop formation and information readout from the genome.

We were able to quantitate the number of genes where Z-flipon variants are causal for mendelian diseases by starting with experimentally validated Z-DNA forming sequences and using these results to predict additional Z-flipons in the genome. We found a direct overlap between mendelian disease-causing variants with predicted (n=372) and experimentally validated (n=124) Z-flipons in 8.6% and 2.9% of OMIM genes(n=4343) respectively (Figure 2D). This conservative approach misses those OMIM genes where the variants impacting Z-flipon biology are not in the region of overlap.

The LOF alleles identified were enriched for frameshifts, with homeobox genes and other transcriptional regulators showing increased susceptibility (Supplemental Data 4). The flipons involved are likely those prone to freeze in the left-handed conformation either due to their length or location in genomic regions of high topological stress, resulting in DNA breaks and error prone repair that increases the frequency of variation. Such events may be prevalent early in development when cell cycles are as short as 3 hours and hypertranscription is prevalent ^39^. Despite the low frequency of their occurrence, the Z-flipon LOF variants may produce mendelian disease more often than more common causes of DNA damage because they induce frameshifts with higher penetrance.

We identified additional LOF variants that overlap Z-flipons in the predicted gnomAD-pLOF collection, but which are not currently associated with mendelian disease (Supplemental Figure 22). Their negative impact may be lessened by alternative splicing, as variants affecting splice sites are more frequent in gnomAD-pLOF ^40^ than we observe with direct OMIM Z-flipon overlaps. Other mechanisms such as transcript destabilization, nonsense-mediated RNA decay and limited or tissue-specific expression could also play a role. Additionally, it is likely that many pLOF variants are somatic rather than germline ^41,42^. Z-flipons also overlap nonsynonymous variants that produce mendelian disease. Around 22% affect arginine codons that contain the Z-prone CG dinucleotide. Yet, there is no evidence that these codons are replaced by the alternative less Z-prone AGG or AGG arginine codons, even though the HBA1 locus clearly demonstrates the possibility of wide-ranging codon replacements in Z-flipon sequences (Supplemental Data 4, Supplemental Figure 10), suggesting that Z-flipon forming sequences are of sufficient biological utility to conserve.

We found that many of the effects of Z-flipons in normal cells likely occur at the level of Z-RNA and involve motifs that have a Z-RNA stem paired with a hairpin loop containing an effector domain. One such example in SMAD1 RNA is characterized by RBM4 binding motifs that promote alternative splicing by suppressing use of splice donor sites. Similar motifs with different effector domains were present in SMAD1, SCARF2 and CACANA1 RNAs. We found examples where disruption of a Z-RNA stem by a SNP allele was associated with the reported GWAS phenotype.

We identified a different motif in which an inverted HR formed a Z-RNA stem by base pairing with another HR. The motif was present in ZNF587B RNA, which has 13 C2H2 (two cysteines, two histidines) ZNF and related proteins that also contain ZNFs and a KRAB domain that suppresses the expression of transposons and viruses by binding to relatively conserved sequences in their genomes. Together this family of proteins constitutes an intracellular form of immunity to protect against such threats ^33^. Here we provide evidence that the system is adaptive.

The HR in these proteins links together adjacent ZNFs ^43^. The sequence has some remarkable properties. In addition to having the propensity to form Z-RNA, the repeat sequence has a strong match to a previously characterized intronic splice enhancer ^34^. Additionally, the HR resembles a recombination recognition sequence (RSS) that is cleaved by RAG1 during immunoglobulin gene rearrangement ^44^. These HR properties suggest multiple mechanisms operating at both the DNA and RNA level for adapting the composition of ZNFs to transposon and viral recombinants that rearrange the conserved binding sites recognized by a ZNF array. At the DNA level, a protein like RAG could create new ZNF arrays through site-specific recombination as occurs in B and T cell receptor genes. We did not find evidence for an increased rate of indels or gene fusions associated with ZNFs in cancer datasets, especially in liver tissues where stellate cells express high levels of RAG1. Noteworthy is the elevated level of missense mutation in some cancer types at positions 9 and 11 of many ZNFs ^35^ adjacent to the HR “ACA” sequence that RAG1 would cleave. DNA site specific recombination between ZNF HPs could operate over longer time periods to diversify ZNF arrays. The recombination events may account for the clusters that are now present on chromosome 19 (Supplementary Figure 9) and for the observation that 179 of 252 degenerate Zinc fingers listed in UNIPROT are found in the KRAB domain containing C2H2 ZNF family.

In contrast, generating variation at the RNA level is a much more rapid process ^45^. While RNA editing recodes ZNF, we did not find nonsynonymous edits that affected the key ZNF nucleic binding sites. Instead, we found evidence supporting the possibility of an adaptive system based on trans-splicing within ZNF gene clusters, possibly by occlusion of HR splice enhancer sites by proteins engaging them as Z-RNA. Such RNA recombination events do not change the specificity of the ZNF but generate new permutations to match a novel transposon or viral rearrangement. Those that enable a cell’s survival likely will be fixed in that cell by epigenetic modifications. Alternatively, they may be fixed by reverse transcription ^45^, possibly using a cleaved HR as a primer to embed the new ZNF combination in an existing ZNF gene.

Interestingly, the unique chromatin structure of C2H2 ZNF clusters reduces recombination of these regions by localizing the H3.3 variant to ZNF containing exons through interactions dependent upon ZNF274 and the ATRX chromatin remodeling complex ^46–48^. At the same time, alternative splicing in this region is favored by the increased levels of H3K36me3 present ^49^. A similar chromatin structure is present at telomeres and also decreases recombination. Interestingly, the same structure is also found at the HBA1 locus ^50,51^. Taken together, the findings raise the possibility that this unique chromatin structure enhances evolutionary adaptation by allowing rapid variation in rates of DNA recombination and RNA processing of the associated genes. The diversity of outcomes produced increases the probability that some individuals will survive when an existential threat emerges. Malaria is one pathogen that drives HBA1 variation ^52^, while alternative telomere maintenance in cancer cells through enhanced recombination of chromosomal ends proves another example of how effective this mechanism can be ^53^.

The results we describe here are consistent with a model where ZNAs localize proteins to a site where they act. With Z-DNA, the chromatin structures and condensates formed can enable approximation of distant regions through loop formation. With Z-RNA, the proteins docked to the effector domains promote specific outcomes. In other cases, Z-RNA binding proteins may occlude sites used for splicing or for interactions with noncoding RNAs. The recognition of left-handed DNA and RNA allows efficient localization of the cellular machinery to active loci and foci where ZNA formation is energized. The process exploits the propensity of short repeat sequences to form alternative nucleic acid structures ^54^. Z-flipons otherwise have low intrinsic informational value but are widely distributed through the genome, opening up a number of possibilities for regulating the readout of genetic information ^55,56^. Through their effects on RNA splicing, editing and expression, Z-flipons can affect a wide range of phenotypes. The work here provides a roadmap for further exploration of the fliponware involved.

## Methods

### Experimental Z-DNA training data

Permanganate/S1 Nuclease Footprinting Z-DNA data contained 41 324 regions with total length of 773 788 bp in human ^10^. The original dataset was filtered for ENCODE blacklisted regions. For DNABERT the data was preprocessed by converting a sequence into 6-mer representation. Each nucleotide position is represented by a k-mer consisting of a current nucleotide and the next 5 nucleotides. The data was split into 5 stratified folds so we could train 5 individual models with 80% of the data and assess precision and recall using the remaining 20%. Due to the large imbalance between positive (Z-DNA) and negative (not Z-DNA) classes we randomly sampled twice as many of the negative class from the Kouzine et al. human data.

### Deep learning transformer-based model training

DNABERT was fine-tuned for the Z-DNA segmentation task with the following hyperparameters: epochs =3, max_learnirng_rate = 1e-5, learning_rate_scheduler = one_cycle (warmup 30%) batch size = 24. We trained 5 models, each on 80% of the positive class examples, and randomly sampled negative class examples. For each 512 bp region from the whole genome the final prediction was made by averaging the predictions of the models that used data not seen during training.

### Model performance

To estimate the model performance we computed precision, recall, F1 and ROC AUC on the test set and for whole-genome predictions (Supplemental Table 1). For benchmark models we applied DeepZ and Gradient boosting methods. DeepZ model was run with the set of 1054 omics features as described in ^8^ for human Shin et al. data set ^9^. Predictions for the test set and whole genome were done the same way as for DNABERT models. CatBoost ^11^ was selected as a gradient boosting benchmark model since CatBoost can use categorical features as an input. The boosting model was trained on the same training set as DNABERT and DeepZ. Each segment from the training set has been encoded into boosting records. Each nucleotide was transformed into DNA segment with 256 + 5 nucleotides. The DNA segment was decomposed in 256 6-mers, and every 6-mer from this DNA segment was mapped to a number from a set of all possible enumerated 6-mers. The resulting categorical vector of length 256 was subsequently used as an input for a boosting model. The Z-DNA was located in the center of the 256 bp DNA targets. All encoded sequences formed a training set that was randomly down sampled to 400 000 objects due to calculation limitations. Test set measurements were performed on the whole test set encoded in the same way.

### Attention visualization

Attention visualization was done with DNABERT-viz tool as described in the original DNABERT paper ^7^.

### Mutagenesis maps

To produce mutagenesis maps, Z-DNABERT was first run using original sequence, then for each site, every nucleotide was replaced with the three alternative nucleotides and the effect of each substitution was calculated as the sum of log(1+p) over each sequence position where p is the probability of Z-DNA formation predicted by the model. By adding 1 to p, we avoided problems with taking the log of a zero probability. The approach allows us to take into account the effects of adjacent sequences on Z-DNA formation, incorporating information of junction formation and cooperativity effects that drive the transition. The heatmap shows the effect of each substitution relative to the original sequence, with the ratio of the two scores reflecting the probability that each will form Z-DNA in that particular context.

### Z-flipon overlap with quantitative trait loci and sites of alternative RNA processing

GWAS catalogue data files were downloaded from https://www.ebi.ac.uk/gwas/ (v. 1.0) ^57^. Data on eQTL, sQTL and edQTL were download from The GTEX portal https://www.gtexportal.org/ (v 8.0) The Swiss Bioinformatics Institute track for alternative splicing ^58^ was accessed through the UCSC browser. Annotation for ENCODE cCREs combined from all cell types was downloaded from UCSC genome browser (data last updated 2020-05-20). Deleterious protein variants were downloaded from the gnomAD-pLOF database (v 2.1.1) ^37^.

### Z-flipon overlap with RNA-editing databases

Z-RNA editing sites from 1413 genes in lung adenocarcinoma tumors was taken from Sharpnack et al. research ^30^. 113 ADAR1 p150-dependent sites were taken from ^29^ Editing sites, associated with Alzheimer’s Disease, were downloaded from ADeditome database ^59^ and also from Rediportal (http://srv00.recas.ba.infn.it/atlas/search.html) ^60^.

### RNA structural analysis

RNA secondary structure was predicted with RNA-fold from Vienna Package ^61^.

### Haplotype Analysis

Haplotypes were determined using the LDLink tool ^62^. Each haplotype was scored by assigning +1 to the alleles that increased trait values and −1 otherwise. For SNPs where quantitative trait measures were unavailable, each allele was assigned a value of 0.

## Availability and implementation

The code is freely available at: https://github.com/mitiau/Z-DNABERT

The Z-DNABERT tool is freely available at:https://colab.research.google.com/github/mitiau/Z-DNABERT/blob/main/ZDNA-prediction.ipynb

## Acknowledgements

The publication was supported by the grant for research centers in the field of AI provided by the Analytical Center for the Government of the Russian Federation (ACRF) in accordance with the agreement on the provision of subsidies (identifier of the agreement 000000D730321P5Q0002) and the agreement with HSE University No. 70-2021-00139.

## Contributions

DU developed Z-DNABERT while DK, AD, NB and AF contributed analyses under the direction of VK, AH and MP. AH and MP wrote the manuscript and prepared figures with assistance from the other co-authors.

## Supplemental Materials

### Z-DNABERT: Fine-tuning DNABERT for Z-DNA prediction

When comparing the test set and whole genome prediction results (Supplemental Table 1), the recall metric does not change much, so the models correctly find all the regions labeled as Z-DNA. Meanwhile, the precision drops sharply, indicating many false positives in the model’s predictions. These false positives could be predictions of novel potential Z-forming regions that were not detected under the experimental conditions used as only a subset of all-possible Z-flipons is active in the cell line used. Supporting this idea is the higher precision of the Kouzine et al data compared to ChIP-seq data. The former has more nucleotides labelled as Z-DNA 0,02% (815 thousand out of 3 billion) compared to the human ChIP-seq data (0,004%, 136 thousand out of 3 billion nucleotides). Also, very high ROC-AUC metrics on whole-genome data show that the model false-positives have probability scores consistently lower than true positives, which could indicate that the regions detected as false-positive are actually regions which have a lower probability of forming Z-DNA in the cells tested.

### Learning Z-DNA sequences from attention maps

It was noticed experimentally that CG/TG/CA repeats are more prone to flip from B- to Z-conformation. However, the detailed analysis of experimentally determined Z-DNA regions showed that other sequences also form Z-DNA, including sequences such as GGGG where the pyrimidine base is replaced by guanine. Transformer architecture allows interpretation of important features by analysis of attention maps. Results can be interpreted according to the difference in the expected frequency of k-mers in the input sequence versus their rank in the output and compared to the frequency in the genome or in the genomic region of interest. This approach is helpful for assessing ChIP-seq data, as a priori, the distribution of ZHUNT3 predicted Z-flipons in the genome is highly biased towards promoters. Many sequences associated with promoters, such as TATA boxes or GC rich segments will have high frequencies in the pull-downs independently of their ability to flip to Z-DNA.

The distributions of 6-mers according to their rank in the attention map are given in Supplemental Table 2. When the model is learning it pays attention not only to the k-mers inside Z-DNA regions but also to the k-mers in the flanking regions. For example, according to attention ranking k-mer GGGGAA is the 7th most frequent that the model uses to define Z-DNA, however this k-mer is the 40th according to the frequency of occurrence inside Z-DNA regions. Also, k-mers GGGGAA CAGGGA TGGGGA GGGGGA AGGGAG GGGAGC are rarely at the site of Z-DNA nucleation, they likely can propagate the flip to Z-DNA once it is initiated. In the model, they appear important for Z-DNA prediction in the nearby sites where alternating pyrimidine/purine sequences may be the first segments to flip conformation.

To investigate further how Z-DNABERT recognizes GT and CA repeats we selected regions from Kouzine et al. human dataset. These repeats are located within 10 bp of each other. Summary attention heatmaps (sum of attention weights from all 12 heads for each position) for various regions are depicted in Supplemental Figure 1.

### Z-DNABERT cross-species predictions and other applications

We tested how well Z-DNABERT model trained on one genome can predict Z-DNA regions in another genome. Supplemental Figure 3 show the results of the model performance that was trained on mouse and then applied on human genome using Kouzine et al. data sets. Performance metrics remain high. The Z-DNA prediction tool published at https://colab.research.google.com/github/mitiau/Z-DNABERT/blob/main/ZDNA-prediction.ipynb allows a user to input sequence into our pretrained model to identify Z-flipons with a high level of confidence.

