## Supplemental Tables for "Z-Flipon Variants reveal the many roles of Z-DNA and Z-RNA in health and disease"

**Supplemental Table 1.** Comparison of Z-DNABERT with other Z-DNA prediction ML models. Predictions are made both on validation set and at the genome-wide level.

| **Validations on the test set:** | | **Precision** | **Recall** | **F1** | **ROC AUC** |
|---|---|---|---|---|---|
| **Human Shin et al.** *Zaa ChIP-seq on HeLa cells* | DeepZ | 0.59 | 0.56 | 0.57 | 0.94 |
| | Z-DNABERT | 0.68 | 0.43 | 0.53 | 0.95 |
| | CatBoost | 0,70 | 0.27 | 0.39 | 0.92 |
| **Human Kouzine et al** | DeepZ* | 0.01 | 0.30 | 0.023 | 0.893 |
| | Z-DNABERT | **0.78** | **0.89** | **0.83** | **0.99** |
| | CatBoost | 0.01 | 0.17 | 0.03 | 0.98 |
| Whole-genome predictions | | **Prec@0.5** | **Recall@0.5** | **F1@0.5** | **ROC AUC** |
| **Human Shin et al.** *Zaa ChIP-seq on HeLa cells* | DeepZ | 0.11 | 0.27 | 0.16 | 0.92 |
| | Z-DNABERT | 0.01 | 0.48 | 0.02 | 0.95 |
| | CatBoost | 0.06 | 0.29 | 0.10 | 0.91 |
| **Human Kouzine et al** | DeepZ* | 0.04 | 0.01 | 0.02 | 0.88 |
| | Z-DNABERT | 0.12 | 0.73 | **0.20** | **1.00** |
| | CatBoost | 0.02 | 0.18 | 0.03 | 0.97 |

**DeepZ that was trained on Shin et al. data*

**Supplemental Table 2.** The top 21 6-mers: Z-DNABERT attention rank versus the 6-mer frequency rank in the experimental datasets tested for tuning the model. The model based on the nucleotide resolution Kouzine et al. data was used in the paper rather than the much smaller 150 basepair resolution ChIP-seq data of Shin et al.

| | hg38 Kouzine et al | | hg38 Shin et al | |
|---|---|---|---|---|
| Attention Rank | 6-mer | Frequency | 6-mer | Frequency |
| 1 | GCGCGC | 1 | TGTGTG | 1 |
| 2 | GTGTGT | 5 | GTGTGT | 2 |
| 3 | CGCGCG | 2 | CGCGCG | 4 |
| 4 | ACACAC | 6 | GCGCGC | 3 |
| 5 | TGTGTG | 3 | CACACA | 5 |
| 6 | GCGCGG | 7 | ACACAC | 6 |
| 7 | CACACA | 4 | ***GGGGAA*** | 40 |
| 8 | CCGCGC | 10 | ***AAAAAA*** | 17 |
| 9 | ***GGG***CGC | 11 | CA***GGG***A | 43 |
| 10 | GCG***CCC*** | 12 | GTGCGC | 11 |
| 11 | GTGCGC | 17 | T***GGGG***A | 331 |
| 12 | GGCGCG | 9 | ***GGGGG***A | 39 |
| 13 | GTGTGC | 14 | GCTGGG | 9 |
| 14 | GCGCAC | 19 | GTGTGC | 7 |
| 15 | GCACAC | 15 | TGCGCG | 8 |
| 16 | GCCCGC | 20 | TGCATG | 21 |
| 17 | GCGGGC | 16 | GGGAAG | 33 |
| 18 | CGCGCC | 8 | AGGGAG | 429 |
| 19 | GCGTGC | 25 | GGGAGC | 458 |
| 20 | GCACGC | 26 | AGAAAG | 38 |
| 21 | ***CCC***GCG | 18 | ***GGGAAA*** | 80 |

**Supplemental Table 3.** DNABERT cross-species predictions

| Trained | Predict | Prec | Recall | F1 | ROC AUC |
|---|---|---|---|---|---|
| **Human Kouzine et al** | hg Kouzine et al. | **0.78** | **0.89** | **0.83** | **1.00** |
| **Mouse Kouzine et al** | hg Kouzine et al. | **0.70** | **0.87** | **0.77** | **1.00** |

**Supplemental Table 4.** Genomic features of predicted and experimental Z-flipons.

| | **Z-flipons** | | | |
|---|---|---|---|---|
| | **Predicted only** | **Experimental and Predicted** | **Experimental only** | **Overlap of Experimental with Predicted** |
| **Promoter (<=3kb)** | 91735 | 26751 | 2040 | 92.91% |
| **5' UTR** | 478 | 90 | 9 | 90.91% |
| **Exons** | 5475 | 1650 | 81 | 95.32% |
| **Introns** | 73807 | 5501 | 527 | 91.26% |
| **3' UTR** | 1641 | 325 | 23 | 93.39% |
| **Downstream (<=300)** | 234 | 32 | 2 | 94.12% |
| **Distal Intergenic** | 76299 | 5810 | 651 | 89.92% |

**Supplemental Table 5.** Direct overlap of ADeditome edits with predicted Z-flipons.

| Location (ADeditome) | ADeditome | ADeditome and Z-DNABERT | ADeditome Percent |
|---|---|---|---|
| downstream | 1805 | 1 | 0.11% |
| exonic | 3686 | 5 | 0.22% |
| exonic;splicing | 9 | | 0.00% |
| intergenic | 17501 | 4 | 1.04% |
| intronic | 1413894 | 226 | 84.34% |
| ncRNA_exonic | 23022 | 3 | 1.37% |
| ncRNA_exonic;splicing | 37 | | 0.00% |
| ncRNA_intronic | 123445 | 18 | 7.36% |
| ncRNA_splicing | 72 | | 0.00% |
| splicing | 255 | | 0.02% |
| upstream | 618 | | 0.04% |
| upstream;downstream | 111 | | 0.01% |
| UTR3 | 87697 | 9 | 5.23% |
| UTR5 | 4116 | 5 | 0.25% |
| UTR5;UTR3 | 95 | | 0.01% |
| Total | 1676363 | 271 | 100% |

**Supplemental Table 6.** ADeditome Genes where predicted Z-flipons are within 1 kb of an Edit.

| | ADeditome Edited Genes | Z-flipon (± 1kb of editing site) | % |
|---|---|---|---|
| Gene Count | **14288** | **6552** | 45.86% |
