## Supplemental Figures for "Z-Flipon Variants reveal the many roles of Z-DNA and Z-RNA in health and disease"

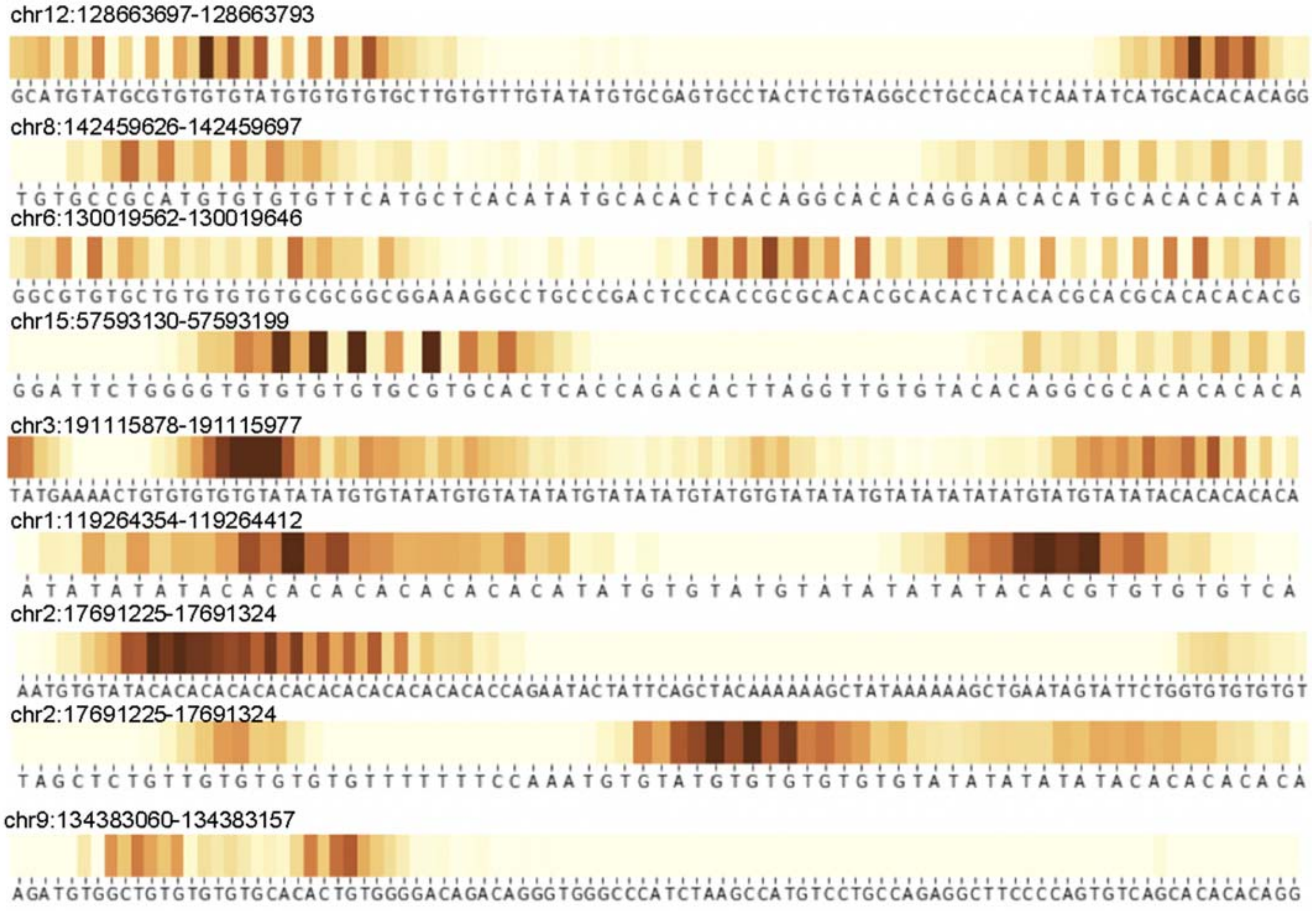


**Supplemental Figure 1.** Samples of attention heatmaps for regions containing both GT and CA repeats in Kouzine et al data set (Kouzine et al., 2017). The dark colors correspond to maximum summary attention. The heaviest stripes at adenosine and guanosine indicates their importance in predicting Z-flipons. The pattern observed is consistent with the role of purines in adopting a *syn* conformation to produce the characteristic *syn-anti* alternation of bases found in Z-DNA.

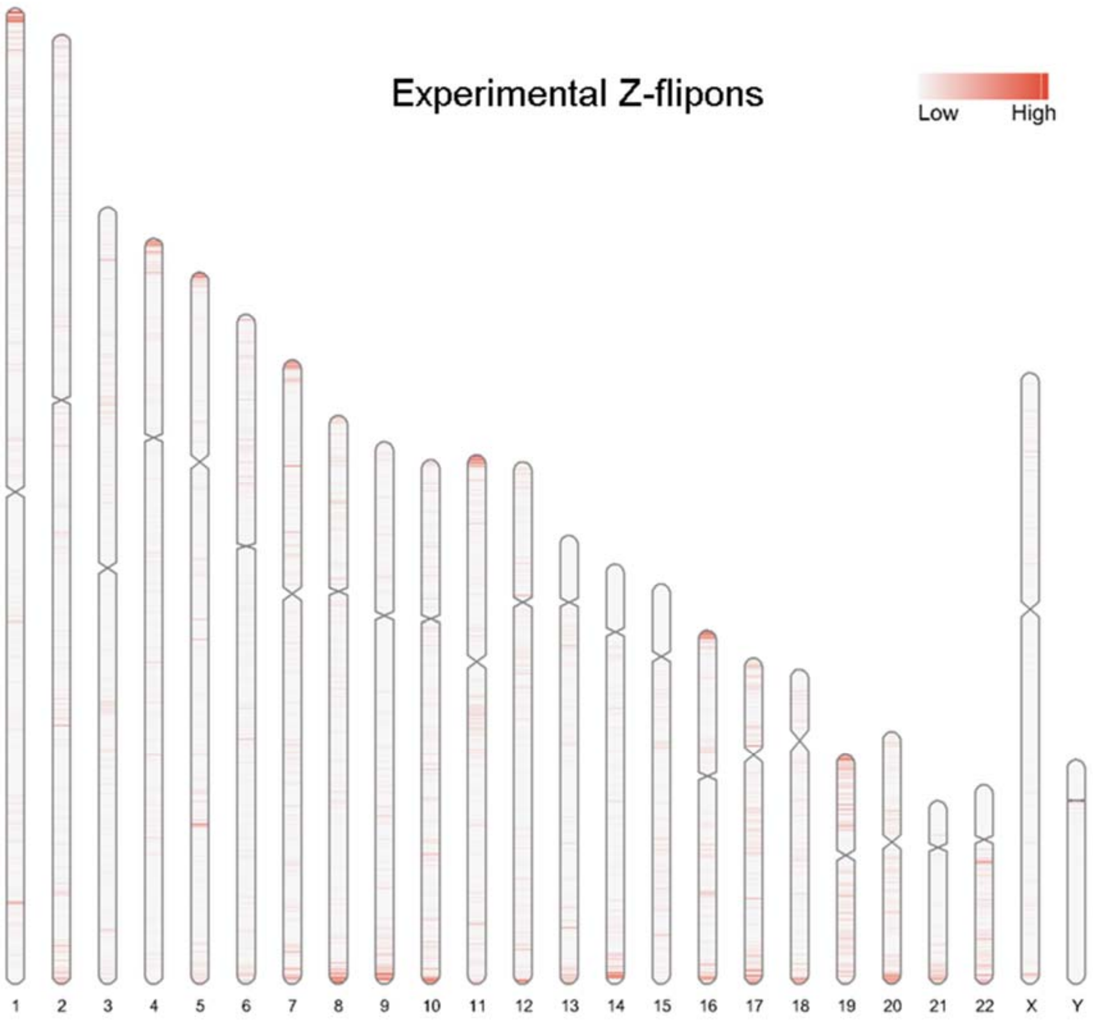


**Supplemental Figure 2**: Ideogram for the Kouzine et al. (Kouzine *et al.*, 2017) experimental Z-DNA data

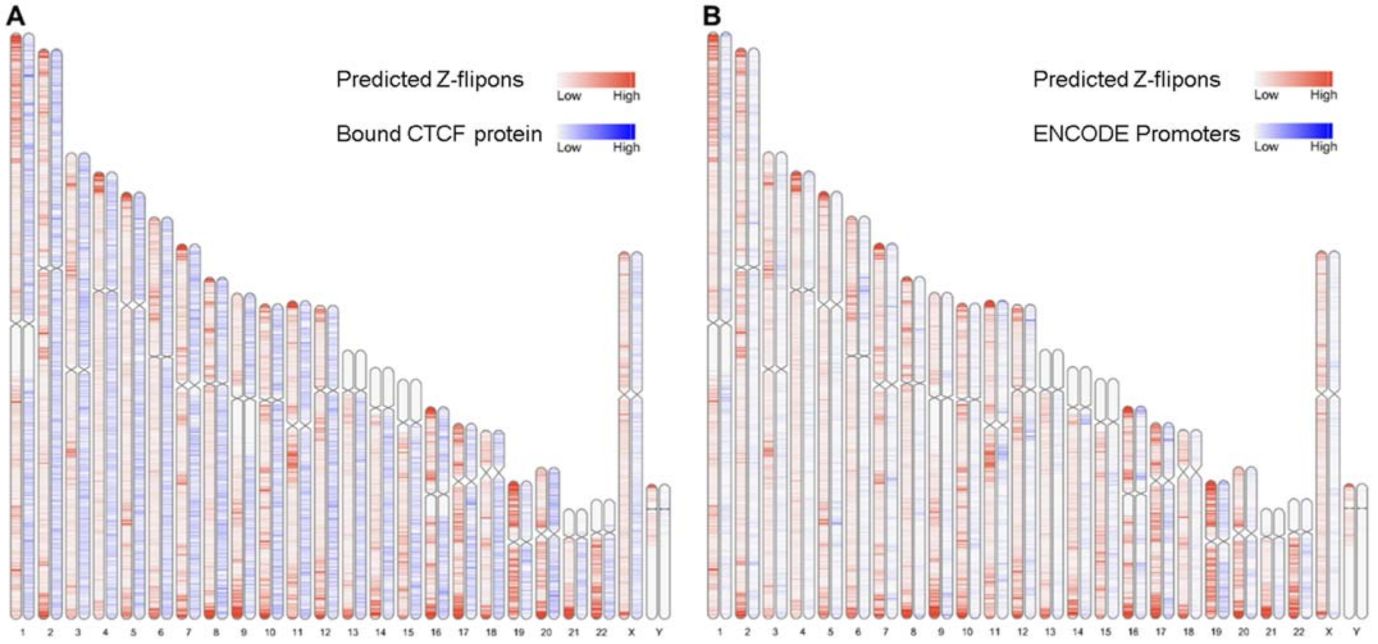


**Supplemental Figure 3** cCRE features from the ENCODE annotations binned in 10 kb windows. **A.** CTCF protein sites **B.** cCRE promoters.

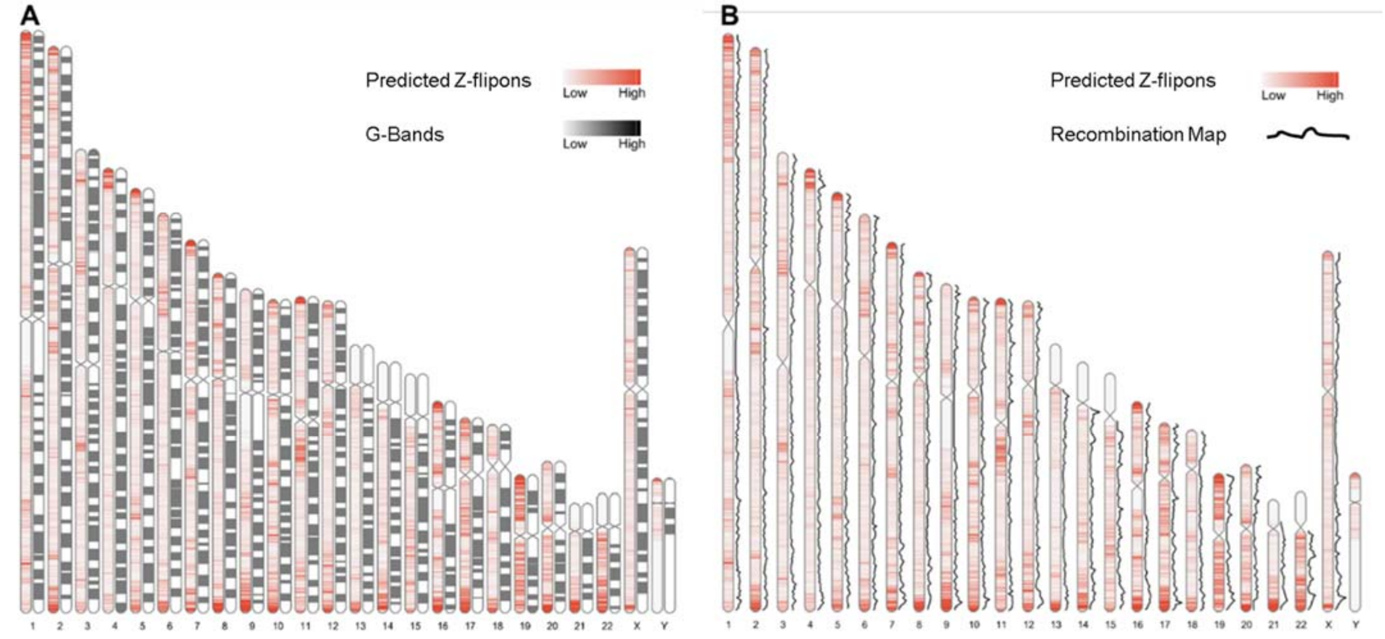


**Supplemental Figure 4.** Z-flipons and Classical Genetic Maps **A.** Location of predicted Z-flipons compared with the chromosomal distribution of G-bands (n=350, shown in black). **B.** The deCode recombination map shown as a back line adjacent to the Z-DNABERT ideogram shown in red. The deCode map is based on 5,136 microsatellite markers for 146 families with a total of 1,257 meiotic events (Kong et al., 2002).

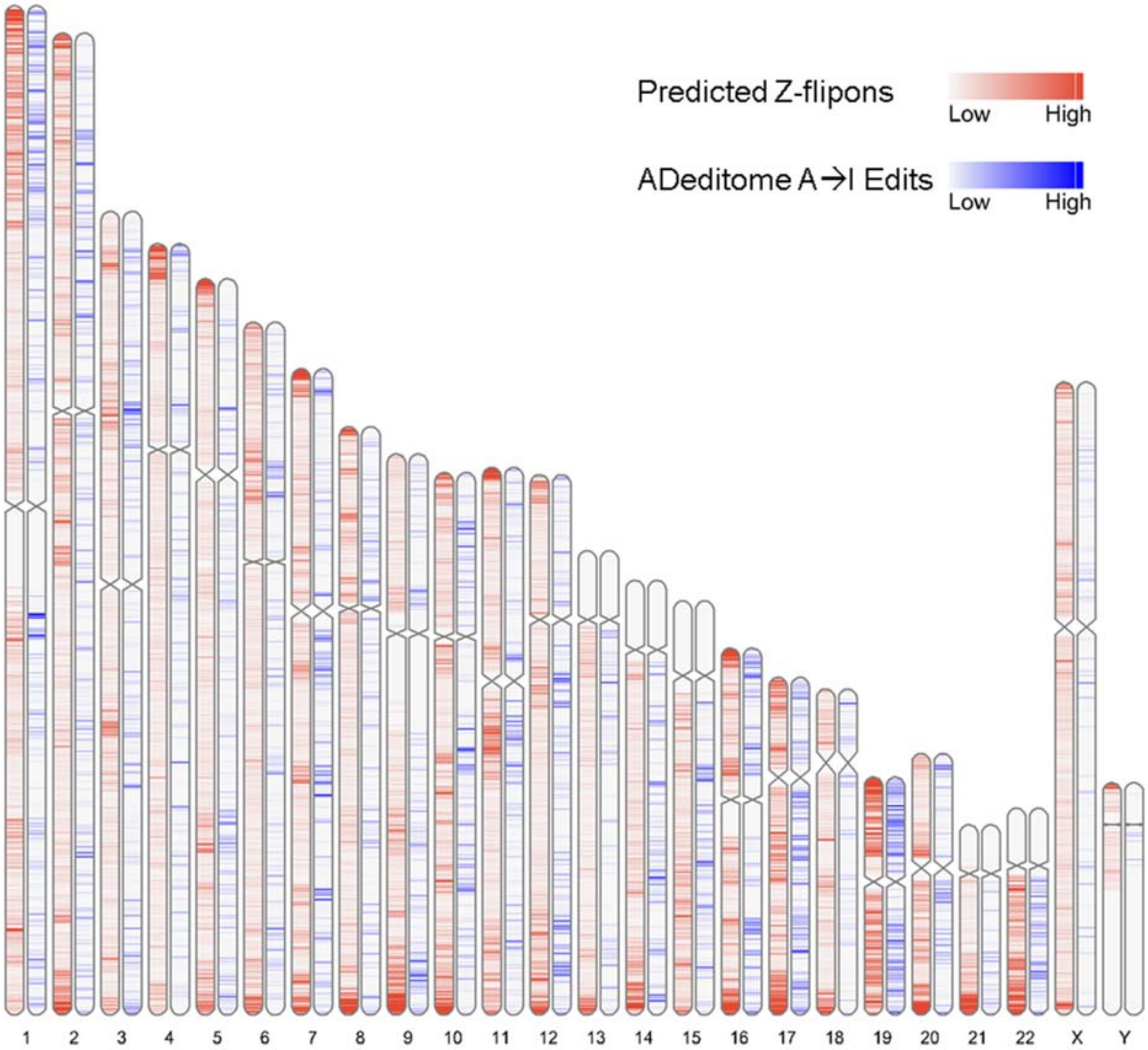


**Supplemental Figure 5.** Ideograms displaying the reported ADeditome edits (blue) alongside the predicted Z-DNABERT regions (red)

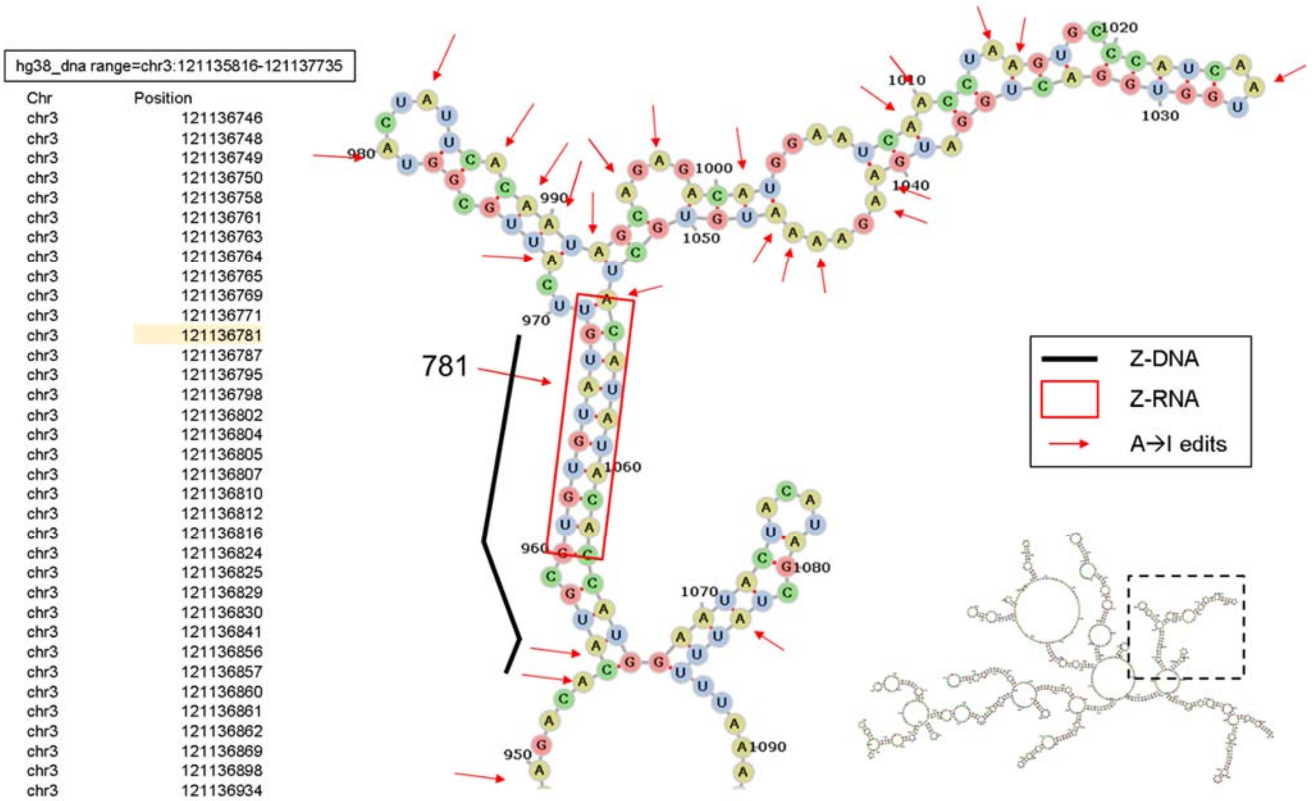


**Supplemental Figure 6**. Example of the STXBP5L intronic RNA fold associated with A→I RNA editing. This RNA is one of the few substrates with an overlap between predicted and experimental Z-flipons and an ADeditome editing site. The Z-flipon detected in the region is indicated by the black line and the Z-RNA forming sequence is in the red box. The editing sites are indicated by red arrows with their genomic position given in the text box on the left. The regional RNA fold is shown at the lower right of the panel with the dotted box indicating the location of the RNA enlarged in the figure. The editing substrate has a Z-stem with the edited adenosines in the associated hairpin loop. The detection of Z-DNA by both computationally and experimental approaches is favored by the 17 basepair length of the Z-RNA stem.

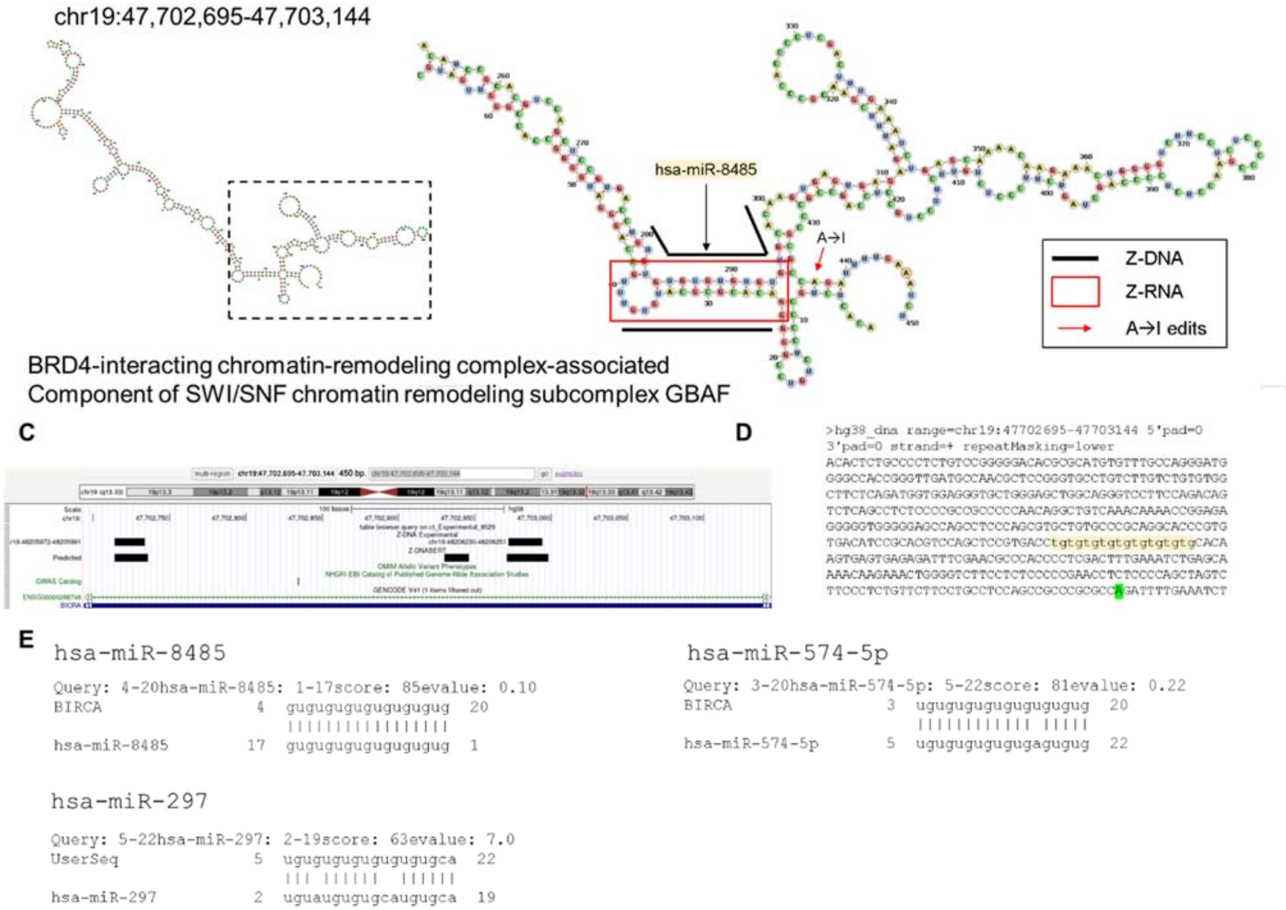


**Supplemental Figure 7**. BICRA (GLTSCR1)( BRD4-interacting chromatin-remodeling complex-associated) that is a component of the SWI/SNF chromatin remodeling subcomplex GBAF. **A.** The RNA fold for BIRCA 3'UTR. **B.** The enlarged view of the boxed region in panel **A**. The fold aligns the Z-RNA stem with an editing site that is documented in multiple databases (seeSupplemental Data 4, lung adenocarcinoma(Experimental)). The black line indicates the position of the experimentally validated Z-DNA **C.** The location of Z-flipons in BICRA 3'UTR. **D.** The sequence recognized by hsa-miR-8485 E. Sequence alignments between BICRA and hsa-miR-8485, hsa-miR-574-5p and hsa-miR-297 (from https://www.mirbase.org/search.shtml).

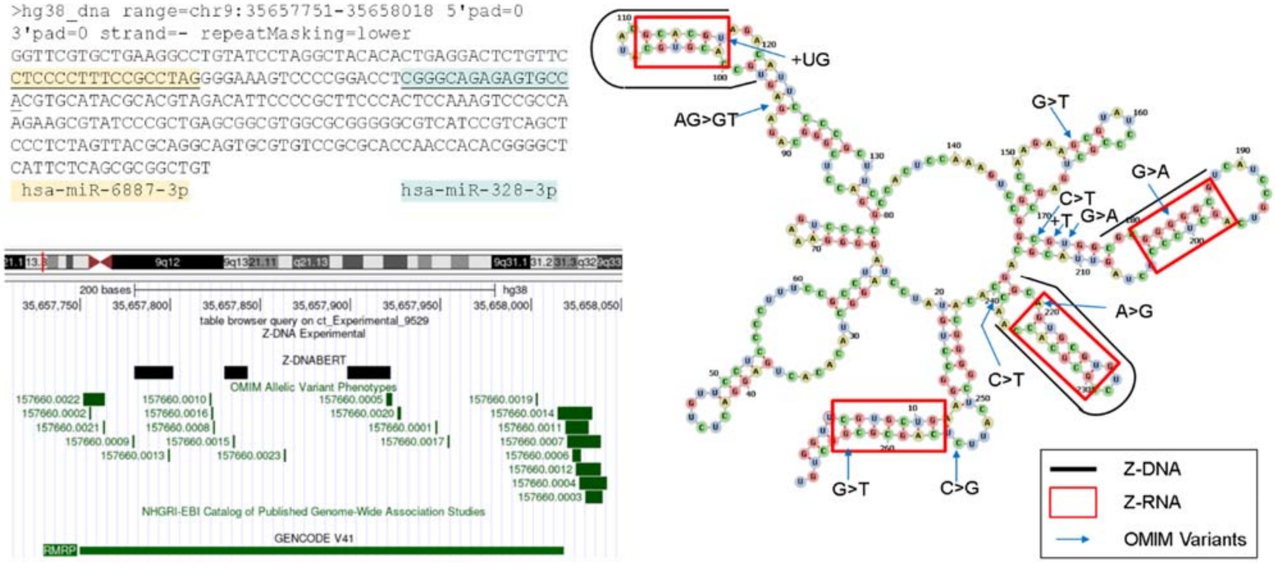


**Supplemental Figure 8**. RMRP noncoding RNA (RNA component of mitochondrial RNA processing endoribonuclease). The RNA formed by folding the RNA is boxed in red with the lines over the seqeunce indicating the regions of Z-DNA predicted by Z-DNABERT. The disease associated variants in OMIM are indecated with an arrow and the reported variation. The colors in the test box indicate the sequences bound by the miRNAs that are listed below the text.

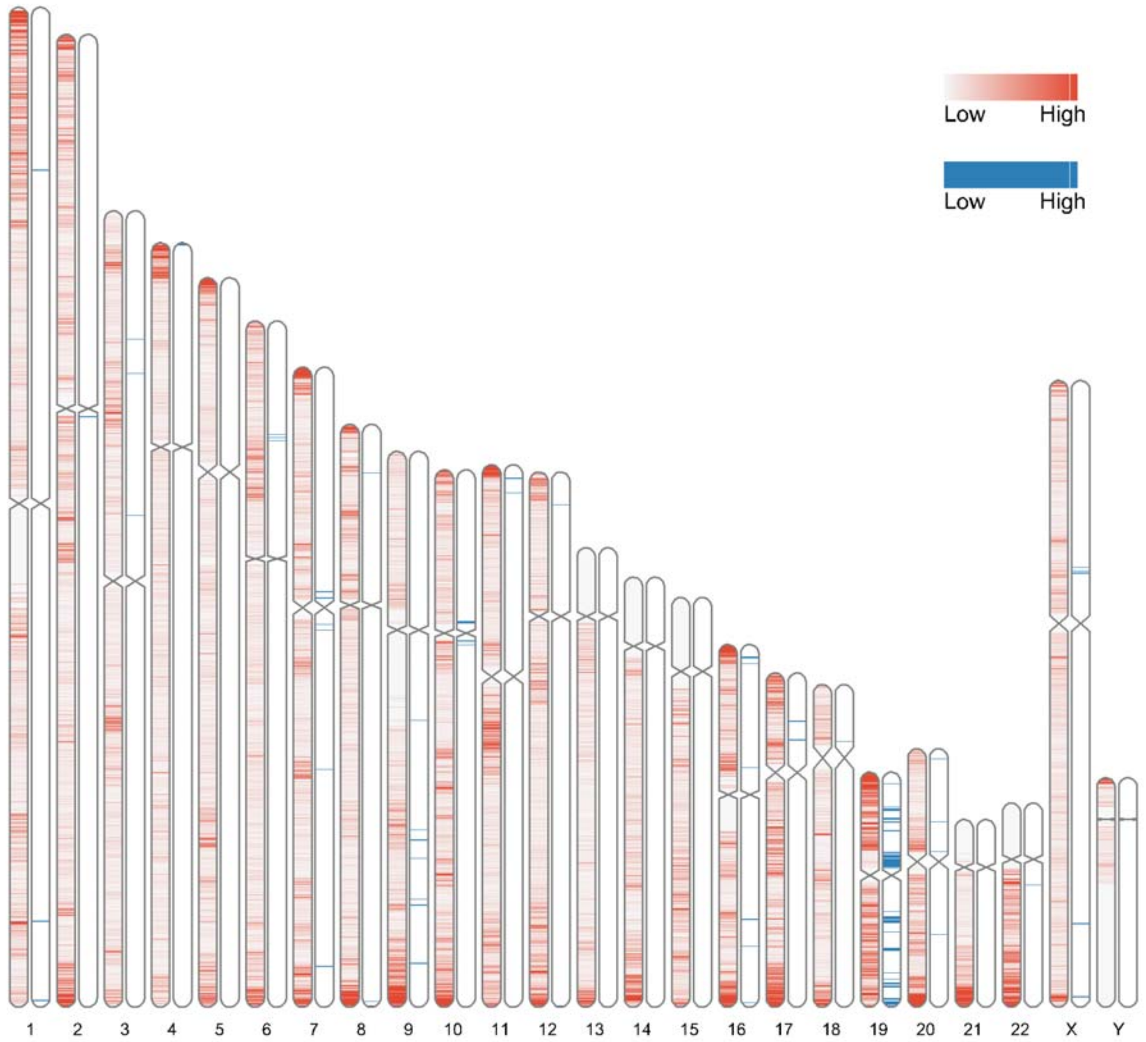


**Supplemental Figure 9**. Ideogram for zinc finger proteins in blue (n=706), with Z-DNABERT predictions in red. Chromosome 19 has many clusters of this protein family.

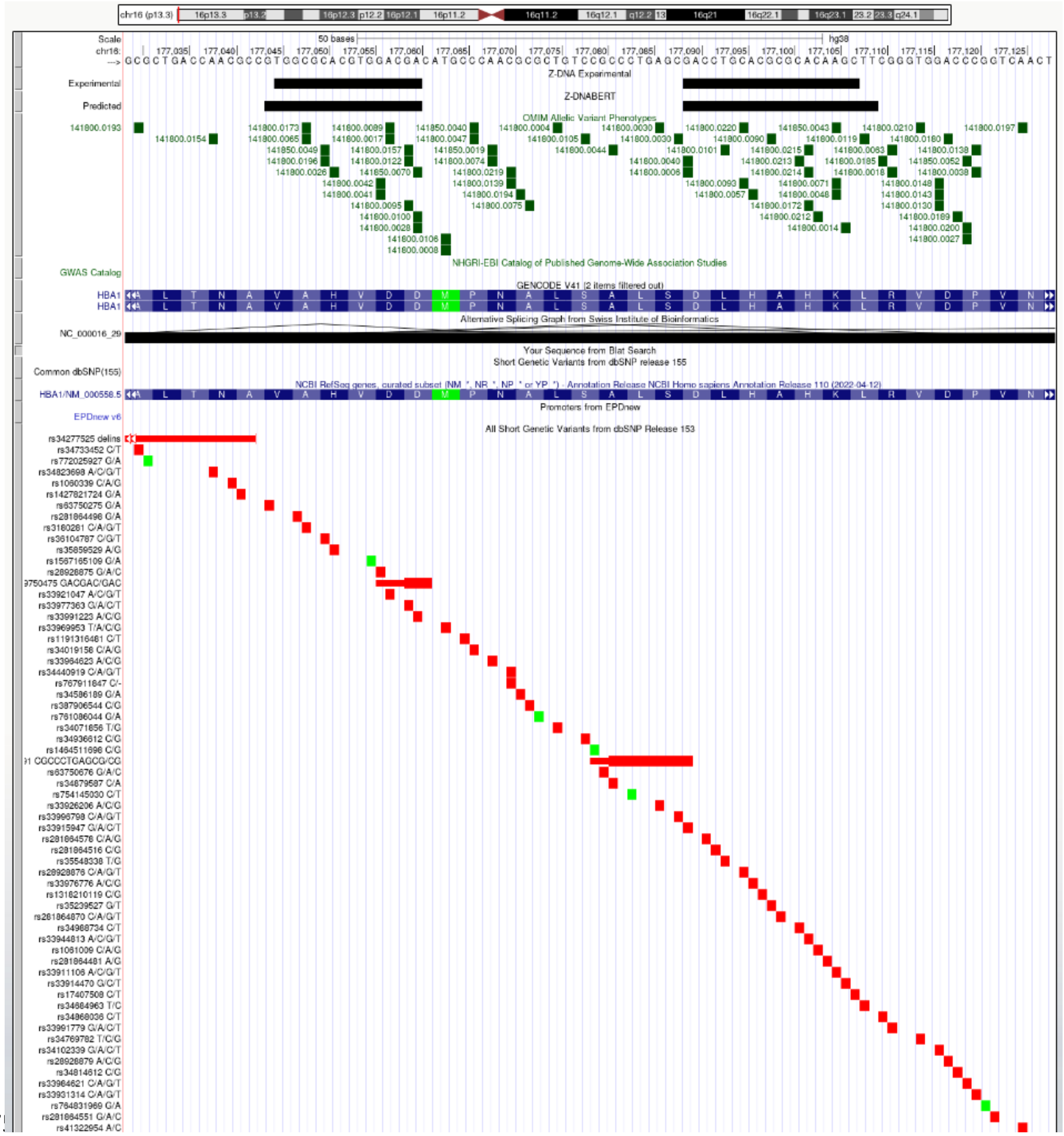


**Supplemental Figure 10. HBA1** (hemoglobin subunit alpha 1). OMIM overlap with predicted and experimental Z-DNA with annotation from the UCSC browser (hg38.chr16:177,029-177,128). Short genetic variants that include predicted gnomAD LOF variants are colored red if the change is nonsynonymous or affects a splice site and green for a synonymous change. Overall, very few gnomAD variants are associated with OMIM phenotypes. The annotation from UCSC for the OMIM variant 141800.0173 is:

OMIM Allelic Variant: 141800.0173 HEMOGLOBIN OZIERI

OMIM: 141800: Hemoglobin alpha-1

Amino Acid Replacement: ALA71VAL

dbSNP/ClinVar: rs3180281

Position: chr16:177048-177048

Band: 16p13.3

Genomic Size: 1

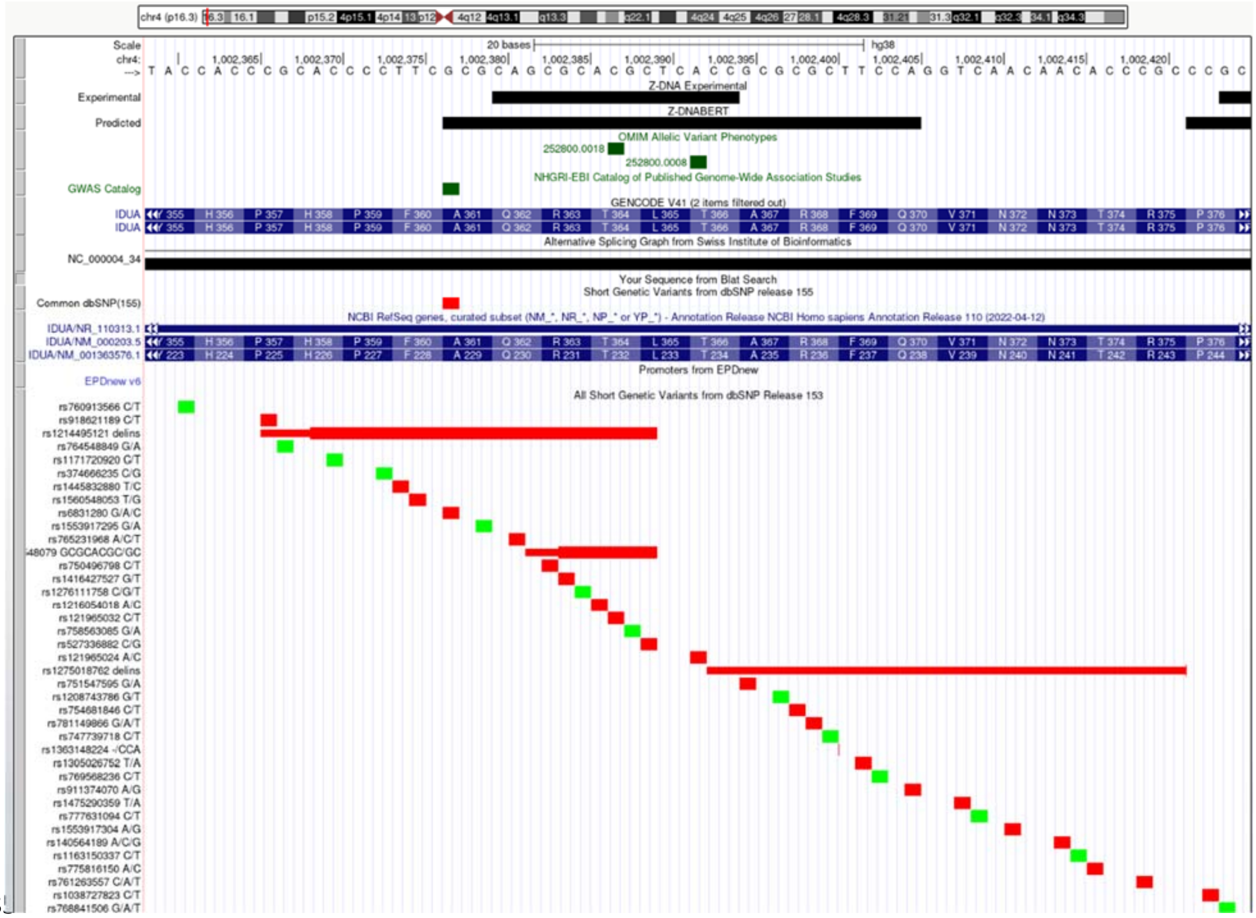


**Supplemental Figure 11.**: **IDUA** (alpha-L-iduronidase). OMIM overlap with predicted and experimental Z-DNA with annotation from the UCSC browser (hg38. chr4:1,002,359-1,002,425). Short genetic variants that include predicted gnomAD LOF variants are colored red if the change is nonsynonymous or affects a splice site and green for a synonymous change. Very few gnomAD variants are associated with OMIM phenotypes. The annotation from UCSC for the OMIM variant 252800.0008 is:

OMIM Allelic Variant: 252800.0008 HURLER SYNDROME

OMIM: 252800: Iduronidase, alpha-L-

Amino Acid Replacement: THR366PRO

dbSNP/ClinVar: rs121965024

Position: chr4:1002392-1002392

Band: 4p16.3

Genomic Size: 1

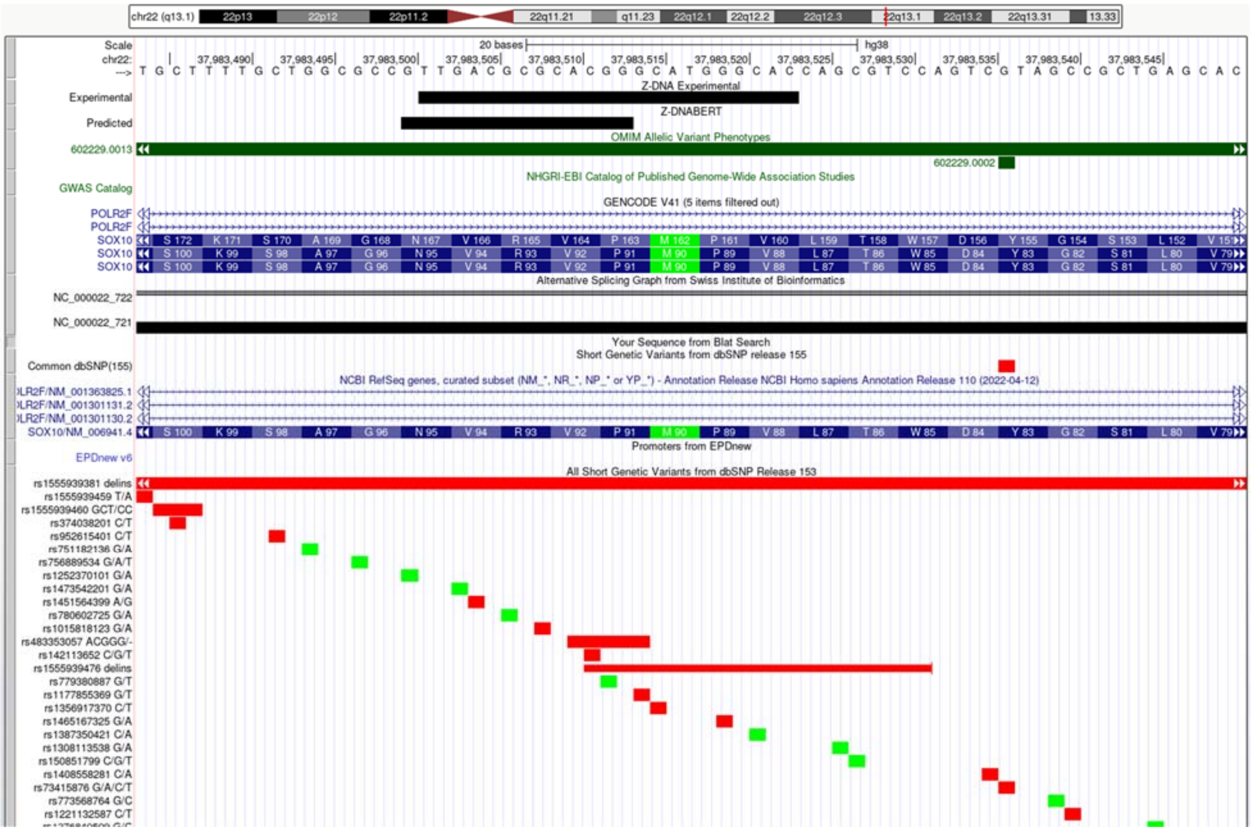


**Supplemental Figure 12.**: **SOX10** (SRY-box transcription factor 10). OMIM overlap with predicted and experimental Z-DNA with annotation from the UCSC browser (hg38. chr22:37,983,484-37,983,550). Short genetic variants that include predicted gnomAD LOF variants are colored red if the change is nonsynonymous or affects a splice site and green for a synonymous change. Very few gnomAD variants are associated with OMIM phenotypes. The annotation from UCSC for the OMIM variant 602229.0013 is:

OMIM Allelic Variant: 602229.0013 WAARDENBURG SYNDROME, TYPE 2E, WITHOUT NEUROLOGIC INVOLVEMENT

OMIM: 602229: SRY (sex-determining region Y)-box-10

Amino Acid Replacement: 253-BP DEL

dbSNP/ClinVar: rs1555939381

Position: chr22:37983314-37983568

Band: 22q13.1

Genomic Size: 255

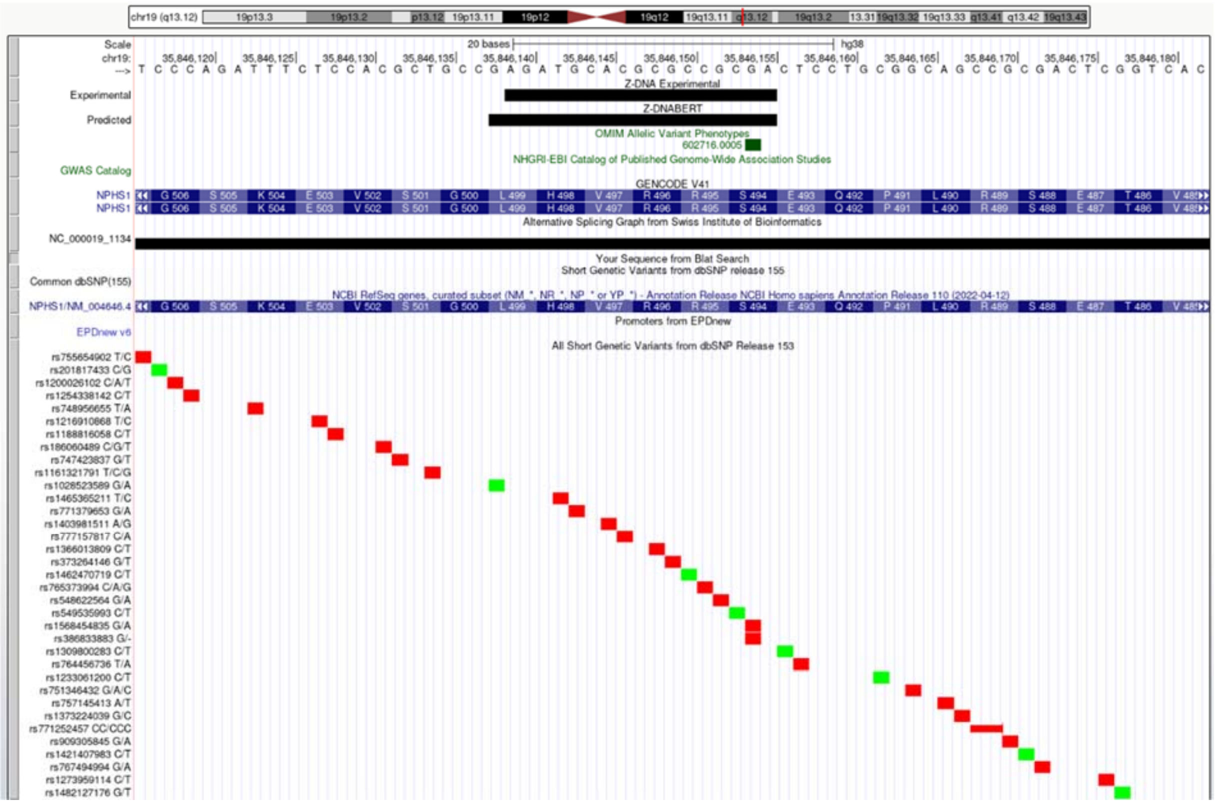


**Supplemental Figure 13. NPHS1** (NPHS1 adhesion molecule, nephrin). OMIM overlap with predicted and experimental Z-DNA with annotation from the UCSC browser (hg38. chr19:35,846,116-35,846,182). Short genetic variants that include predicted gnomAD LOF variants are colored red if the change is nonsynonymous or affects a splice site and green for a synonymous change. Very few gnomAD variants are associated with OMIM phenotypes. The annotation from UCSC for the OMIM variant 602716.0005 is:

OMIM Allelic Variant: 602716.0005 NEPHROTIC SYNDROME, TYPE 1

OMIM: 602716: Nephrin

Amino Acid Replacement: 1-BP DEL, 1481C

dbSNP/ClinVar: rs386833883

Position: chr19:35846154-35846154

Band: 19q13.12

Genomic Size: 1

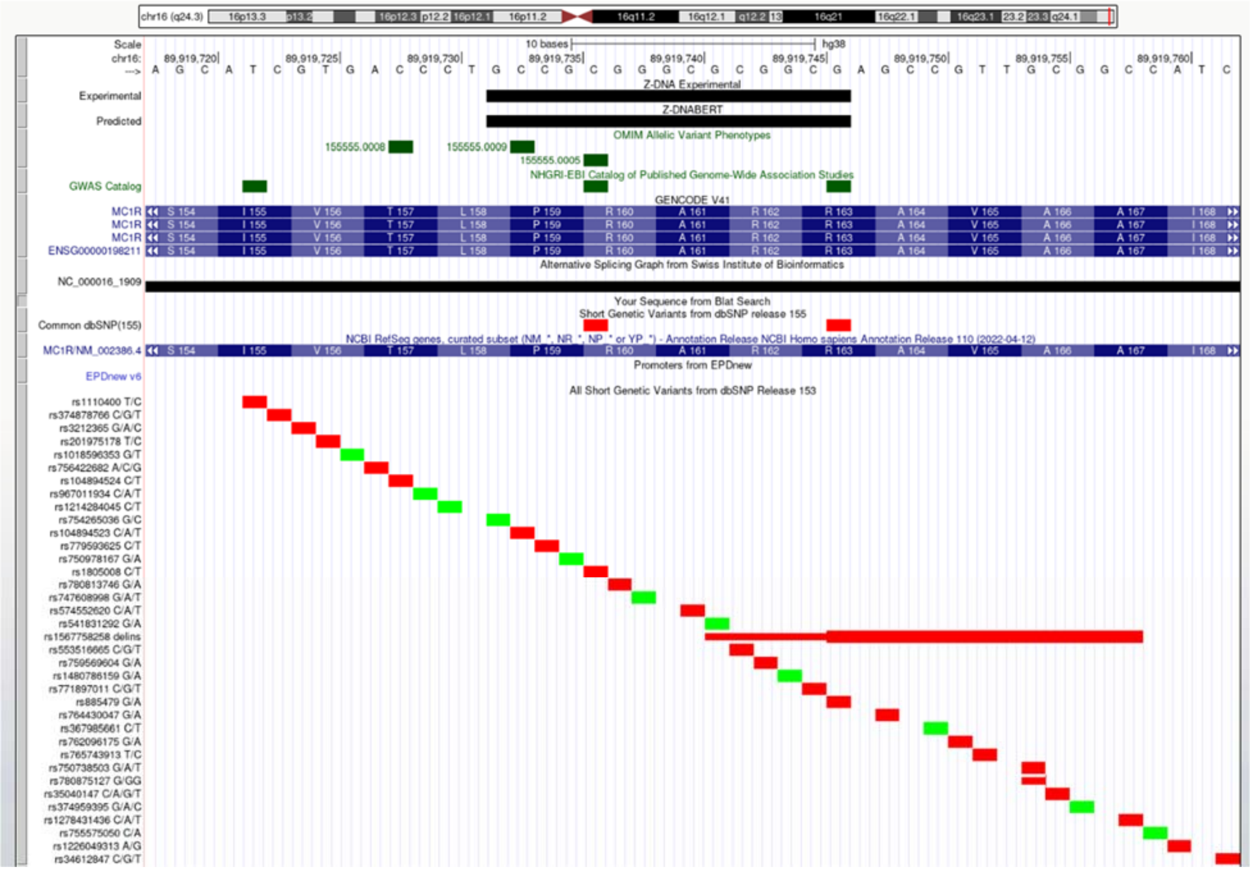


**Supplemental Figure 14. MC1R** (melanocortin 1 receptor). OMIM overlap with predicted and experimental Z-DNA with annotation from the UCSC browser (hg38. chr16:89,919,718-89,919,762). Short genetic variants that include predicted gnomAD LOF variants are colored red if the change is nonsynonymous or affects a splice site and green for a synonymous change. Very few gnomAD variants are associated with OMIM phenotypes. The annotation from UCSC for the OMIM variants 155555.0005 and 155555.0009 are:

OMIM Allelic Variant: 155555.0009 UV-INDUCED SKIN DAMAGE, SUSCEPTIBILITY TO

OMIM: 155555: Melanocortin-1 receptor (alpha melanocyte-stimulating hormone receptor)

Amino Acid Replacement: PRO159THR

dbSNP/ClinVar: rs104894523

Position: chr16:89919733-89919733

Band: 16q24.3

Genomic Size: 1

OMIM Allelic Variant: 155555.0005 SKIN/HAIR/EYE PIGMENTATION 2, RED HAIR/FAIR SKIN

OMIM: 155555: Melanocortin-1 receptor (alpha melanocyte-stimulating hormone receptor)

Amino Acid Replacement: ARG160TRP

dbSNP/ClinVar: rs1805008

Position: chr16:89919736-89919736

Band: 16q24.3

Genomic Size: 1

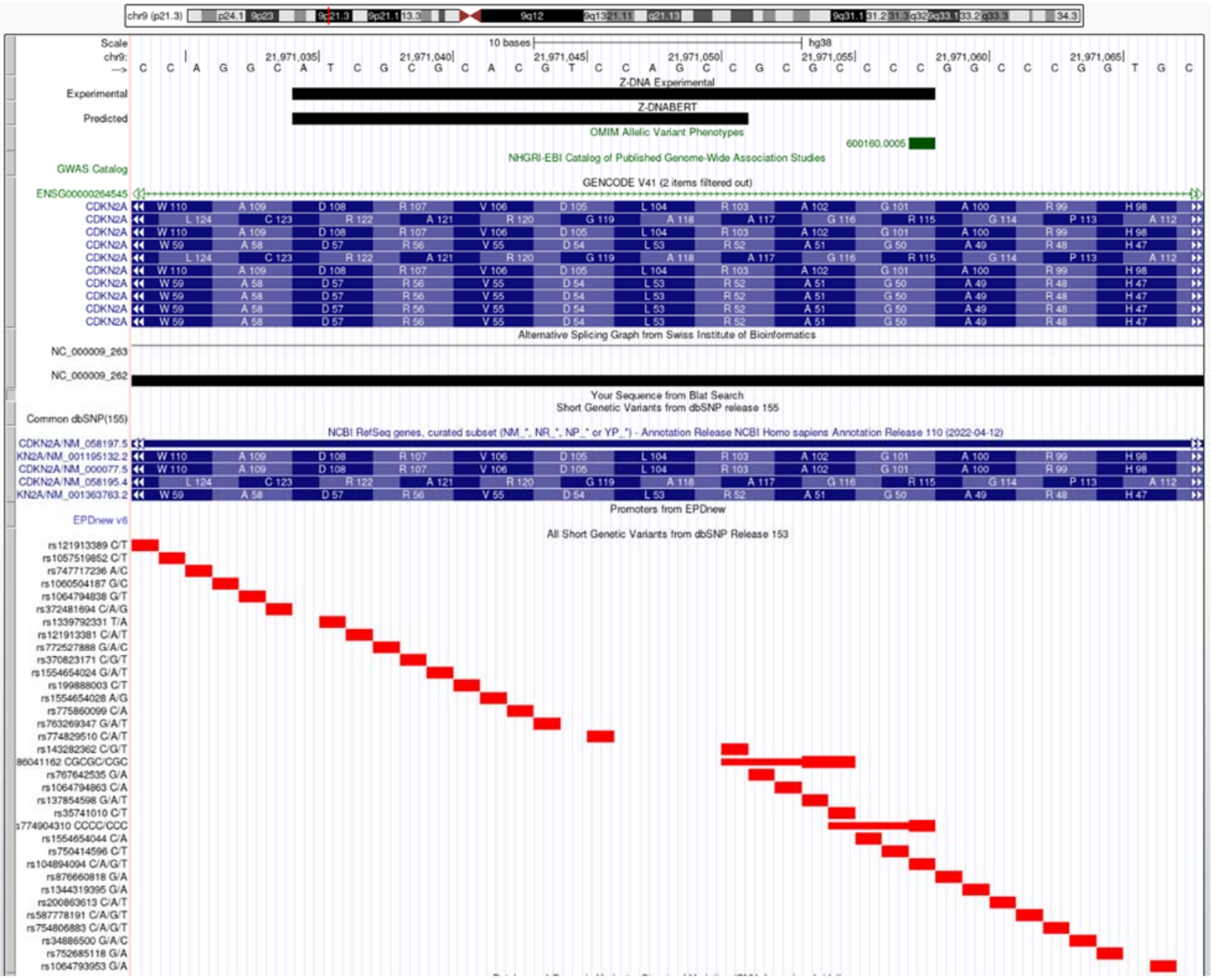


**Supplemental Figure 15.**: **CDKN2A** (cyclin dependent kinase inhibitor 2A). OMIM overlap with predicted and experimental Z-DNA with annotation from the UCSC browser (hg38. chr9:21,971,029-21,971,068). Short genetic variants that include predicted gnomAD LOF variants are colored red if the change is nonsynonymous or affects a splice site and green for a synonymous change. Very few gnomAD variants are associated with OMIM phenotypes. The annotation from UCSC for the OMIM variant 600160.0005 is:

OMIM Allelic Variant: 600160.0005 MELANOMA, CUTANEOUS MALIGNANT, SUSCEPTIBILITY TO, 2

OMIM: 600160: Cyclin-dependent kinase inhibitor 2A (p16, inhibits CDK4)

Amino Acid Replacement: GLY101TRP

dbSNP/ClinVar: rs104894094

Position: chr9:21971058-21971058

Band: 9p21.3

Genomic Size: 1

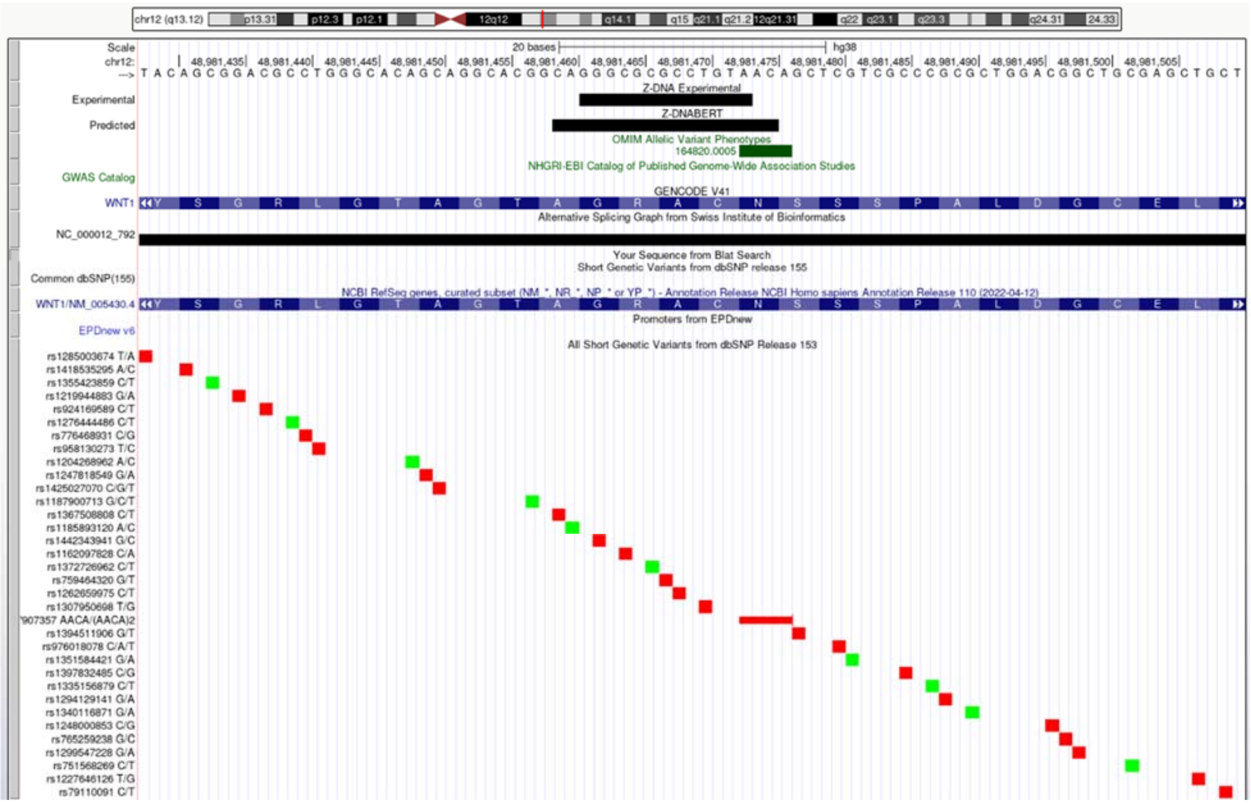


**Supplemental Figure 16.**: **WNT1** (Wnt family member 1). OMIM overlap with predicted and experimental Z-DNA with annotation from the UCSC browser (hg38. chr12:48,981,428-48,981,510). Short genetic variants that include predicted gnomAD LOF variants are colored red if the change is nonsynonymous or affects a splice site and green for a synonymous change. Very few gnomAD variants are associated with OMIM phenotypes. The annotation from UCSC for the OMIM variant 164820.0005 is:

OMIM Allelic Variant: 164820.0005 OSTEOGENESIS IMPERFECTA, TYPE XV

OMIM: 164820: Wingless-type MMTV integration site family, member 1 (oncogene INT1)

Amino Acid Replacement: 4-BP INS, 946AACA

dbSNP/ClinVar: rs387907357

Position: chr12:48981473-48981476

Band: 12q13.12

Genomic Size: 4

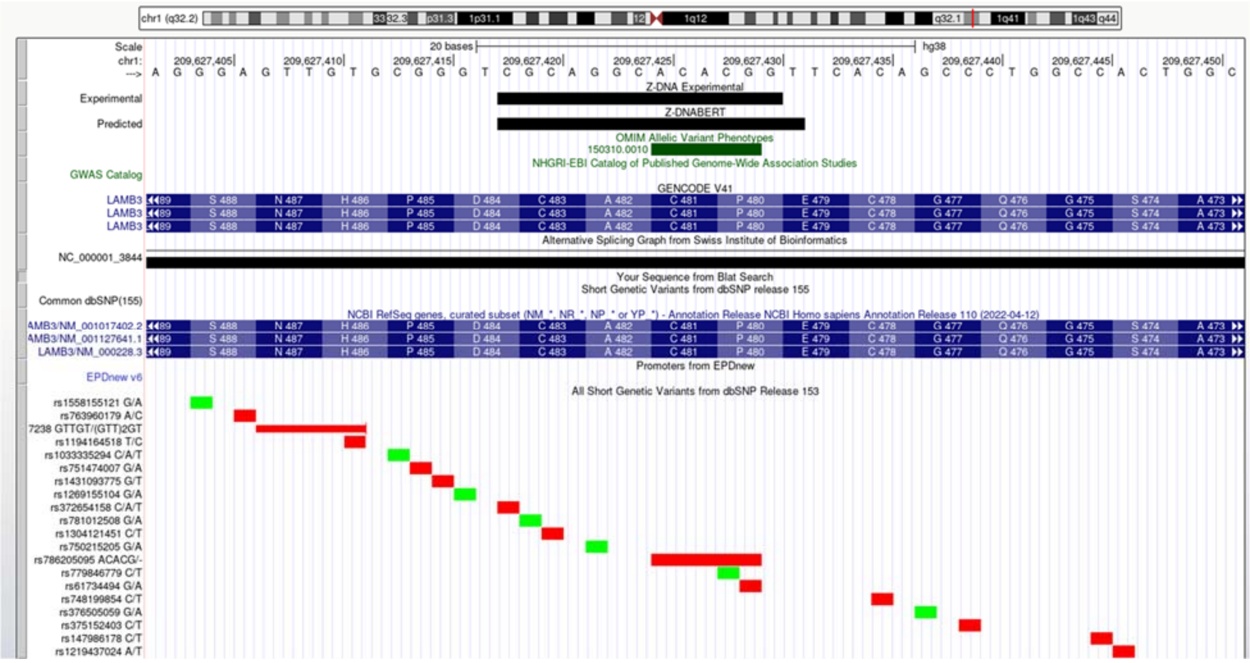


**Supplemental Figure 17.**: LAMB3 (laminin subunit beta 3). OMIM overlap with predicted and experimental Z-DNA with annotation from the UCSC browser (hg38: chr1:209,627,402-209,627,451). Short genetic variants that include gnomAD LOF variants are colored red if the change is nonsynonymous or affects a splice site and green for a synonymous change. Very few gnomAD variants are associated with OMIM phenotypes. The annotation from UCSC for the OMIM variant shown is:

OMIM Allelic Variant: 150310.0010 EPIDERMOLYSIS BULLOSA, JUNCTIONAL 1A, INTERMEDIATE

OMIM: 150310: Laminin, beta-3 (nicein, 125kD; kalinin, 140kD; BM600, 125kD)

Amino Acid Replacement: 5-BP DEL, NT1438

dbSNP/ClinVar: rs786205095

Position: chr1:209627425-209627429

Band: 1q32.2

Genomic Size: 5

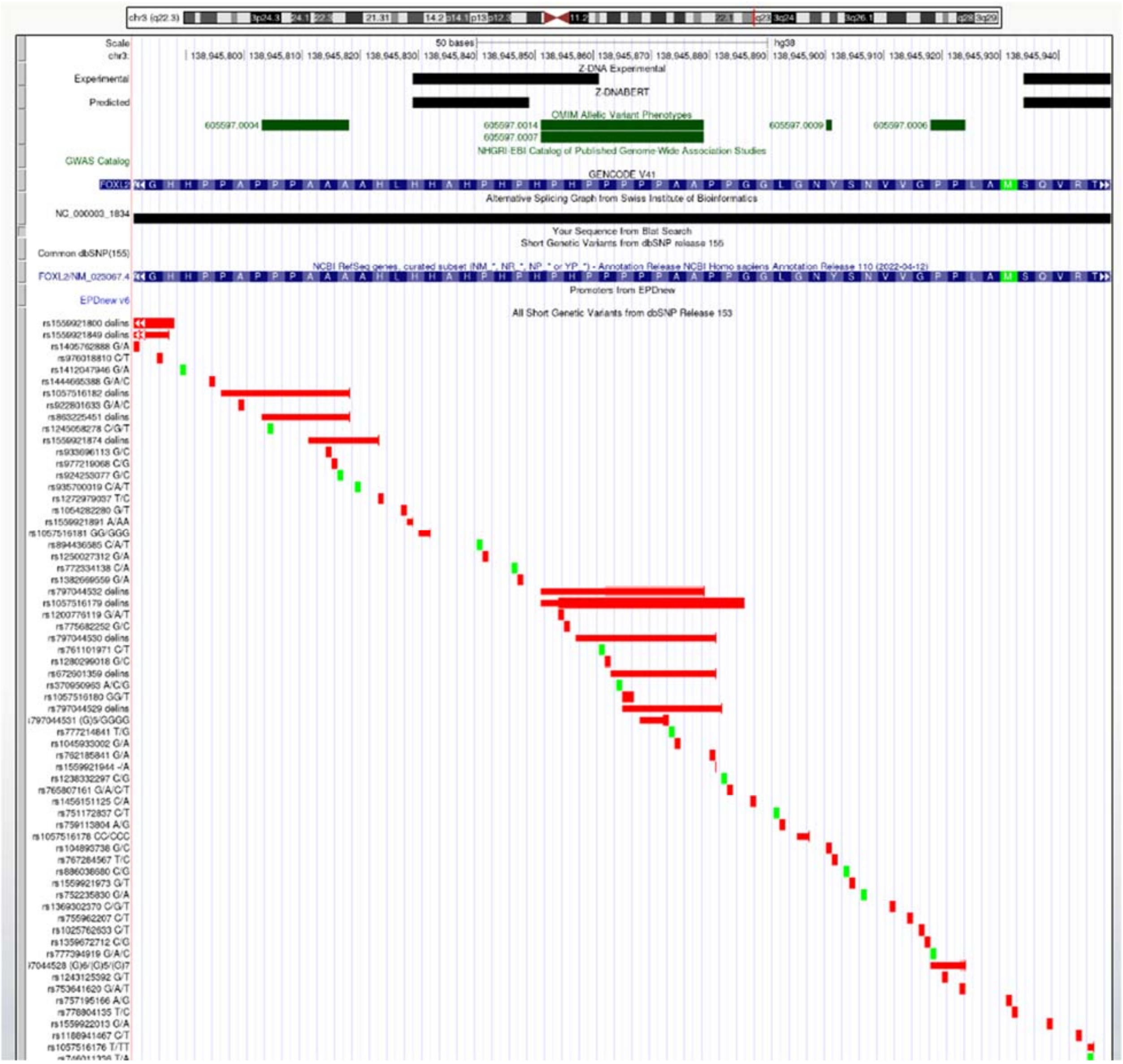


**Supplemental Figure 18. FOXL2** (forkhead box L2). OMIM overlap with predicted and experimental Z-DNA with annotation from the UCSC browser (hg38. chr3:138,945,782-138,945,949). Short genetic variants that include predicted gnomAD LOF variants are colored red if the change is nonsynonymous or affects a splice site and green for a synonymous change. The annotation from UCSC for the OMIM variant 605597.0014 is:

OMIM Allelic Variant: 605597.0014 BLEPHAROPHIMOSIS, PTOSIS, AND EPICANTHUS INVERSUS, TYPE I

OMIM: 605597: Forkhead transcription factor FOXL2

Amino Acid Replacement: 17-BP DUP, NT1092

dbSNP/ClinVar: rs797044532

Position: chr3:138945852-138945879

Band: 3q22.3

Genomic Size: 28

**A.**

| | OMIM NonSynonymous Variants Overlapped by Z-Flipons | | | | | Protein Composition |
|---|---|---|---|---|---|---|
| | Row Labels | Predicted | Percent | Experimetal | Percent | |
| 1 | ALA | 35 | 10.36 | 10 | 8.47 | 8.70 |
| 2 | ARG | 96 | 28.40 | 26 | 22.03 | 6.20 |
| 3 | ASN | 5 | 1.48 | 2 | 1.69 | 3.90 |
| 4 | ASP | 21 | 6.21 | 15 | 12.71 | 5.10 |
| 5 | CYS | 25 | 7.40 | 7 | 5.93 | 1.50 |
| 6 | GLN | 1 | 0.30 | 0 | 0.00 | 3.90 |
| 7 | GLU | 17 | 5.03 | 9 | 7.63 | 6.20 |
| 8 | GLY | 21 | 6.21 | 4 | 3.39 | 6.80 |
| 9 | HIS | 26 | 7.69 | 15 | 12.71 | 2.20 |
| 10 | ILE | 1 | 0.30 | 0 | 0.00 | 5.70 |
| 11 | LEU | 10 | 2.96 | 2 | 1.69 | 9.80 |
| 12 | LYS | 7 | 2.07 | 4 | 3.39 | 5.30 |
| 13 | MET | 7 | 2.07 | 2 | 1.69 | 2.40 |
| 14 | PHE | 2 | 0.59 | 1 | 0.85 | 4.00 |
| 15 | PRO | 10 | 2.96 | 2 | 1.69 | 5.00 |
| 16 | SER | 5 | 1.48 | 2 | 1.69 | 7.00 |
| 17 | THR | 19 | 5.62 | 8 | 6.78 | 5.30 |
| 18 | TRP | 2 | 0.59 | 0 | 0.00 | 1.30 |
| 19 | TYR | 4 | 1.18 | 3 | 2.54 | 2.90 |
| 20 | VAL | 24 | 7.10 | 6 | 5.08 | 6.50 |
| | Grand Total | 338 | 100.00 | 118 | 100.00 | 99.70 |

**B**

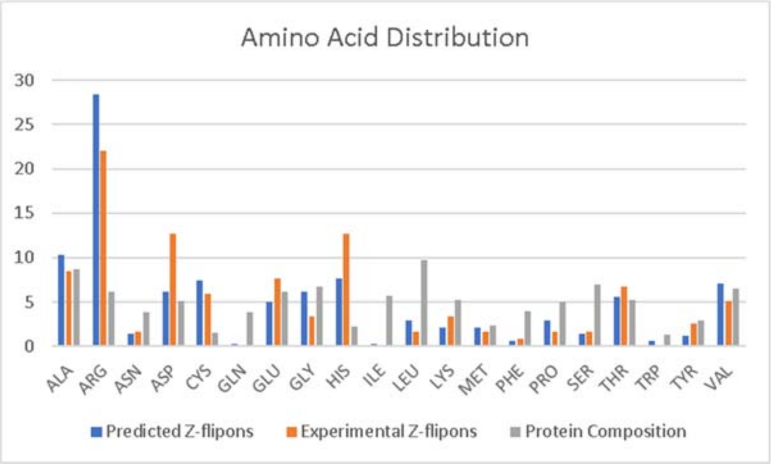


**Supplemental Figure 19. A.** Amino acids altered in OMIM Variants that are overlapped by predicted and experimentally validated Z-flipons. The amino acid frequency in proteins is given for comparison (Trinquier and Sanejouand, 1998). **B.** Graph of Table A.

A Z-Flipon Overlap by OMIM Gene

| | OMIM | Gencode V41 |
|---|---|---|
| | n=4342 | n=19,370 |
| | Percent | Percent |
| experimental (n=124) | 2.86 | 0.64 |
| predicted (n=372) | 8.57 | 1.92 |

B Z-Flipon Overlap by OMIM Variant

| OMIM Variation | Predicted | Percent | Experimental | Percent |
|---|---|---|---|---|
| frameshift_variant | 97 | 19.80 | 36 | 21.56 |
| multiple variants | 3 | 0.61 | 0 | 0.00 |
| ncRNA variant | 4 | 0.82 | 0 | 0.00 |
| nonsynonymous | 337 | 68.78 | 118 | 70.66 |
| splice_acceptor_variant | 2 | 0.41 | 0 | 0.00 |
| splice_donor_variant | 22 | 4.49 | 0 | 0.00 |
| stop_gained | 25 | 5.10 | 13 | 7.78 |
| Grand Total | 490 | 100.00 | 167 | 100.00 |

C

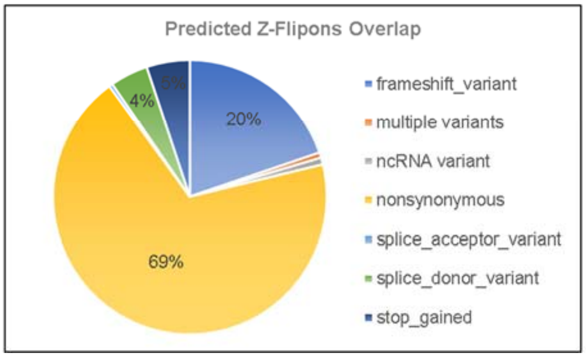


D

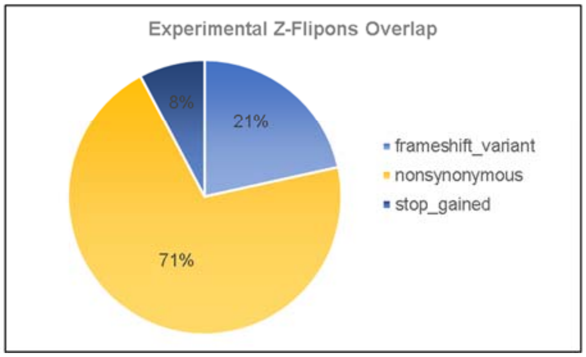


**Supplemental Figure 20**. Flipon overlap with Morbid OMIM **A**. Overlap of Z-flipons and OMIM Genes. **B**. OMIM variants overlapped by experimental and predicted Z-flipons **C**. Variants that overlap predicted Z-flipons **D.** Overlap for experimentally validated flipons.

A

**gnomAD Predicted Loss of Function Variants**

| | Experiment | Percent | Z-DNABERT | Percent | All | Percent |
|---|---|---|---|---|---|---|
| frameshift_variant | 745 | 64.22 | 2543 | 58.30 | 196760 | 44.35 |
| splice_acceptor_variant | 17 | 1.47 | 135 | 3.09 | 45157 | 10.18 |
| splice_donor_variant | 105 | 9.05 | 547 | 12.54 | 56115 | 12.65 |
| stop_gained | 293 | 25.26 | 1137 | 26.07 | 145646 | 32.83 |
| Total | 1160 | 100.00 | 4362 | 100.00 | 443678 | 100.00 |

B

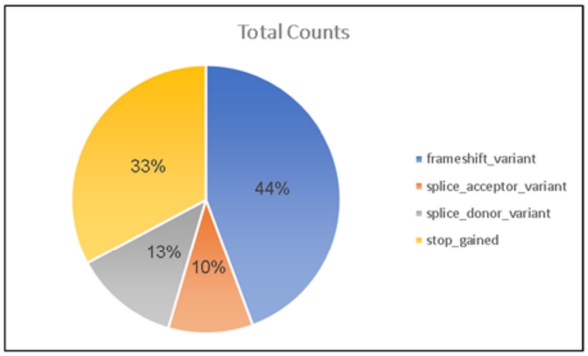


C

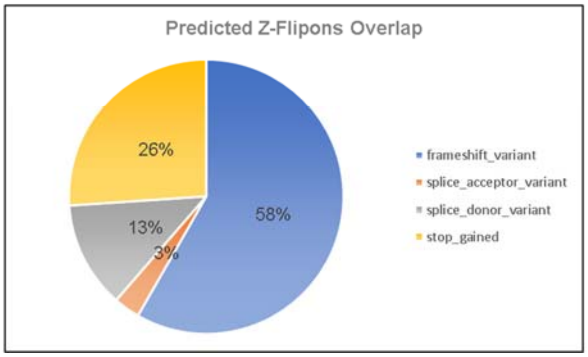


D

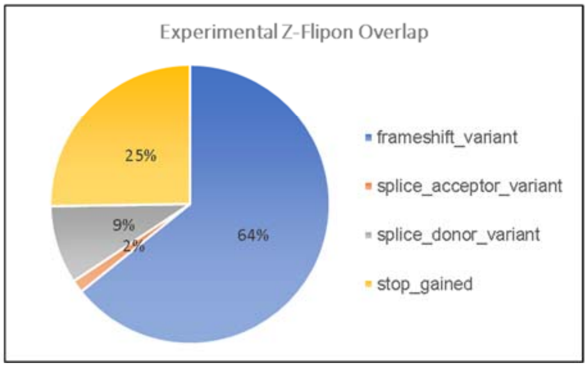


**Supplemental Figure 21**. Overlap of Z-flipons with predicted gnomAD-pLOF variants **A.** Counts **B.** Total predicted gnomAD loss of function variants by class. **C.** Variants in predicted Z-flipons show a higher rate of frameshifts compared to the total collection. **D.** The increase in frameshift variants is also present in the experimentally validated Z-flipons.

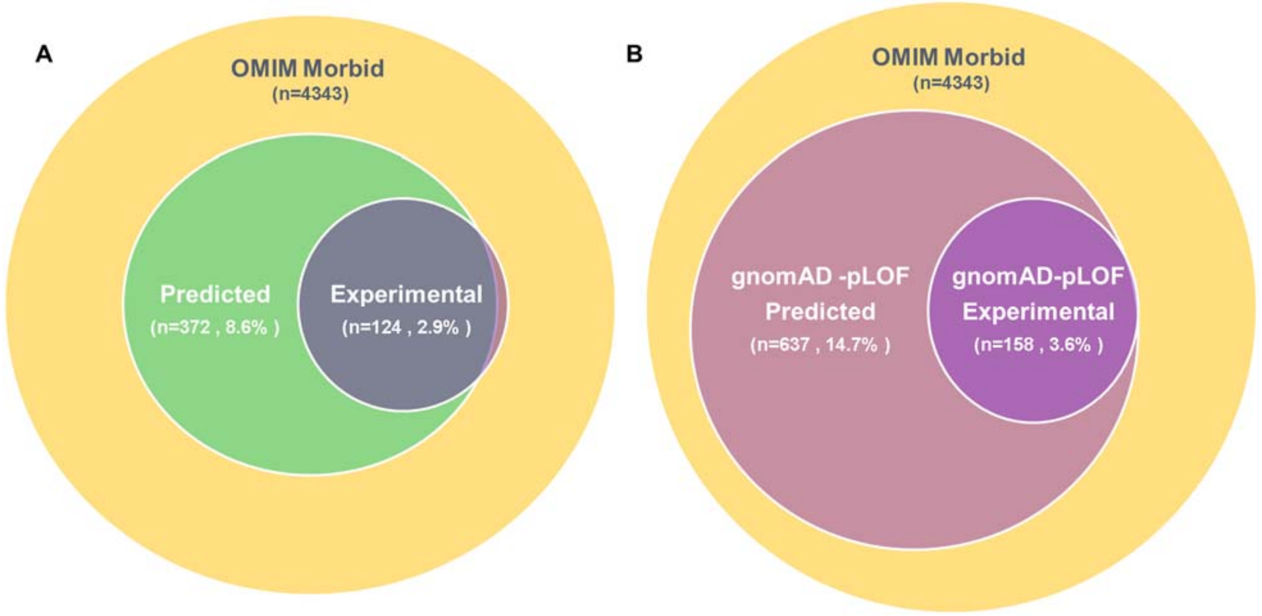


**Supplemental Figure 22.** OMIM Genes **A**. Genes in the Morbid OMIM collection with variants that overlap predicted and experimenta Z-flipons. **B.** Genes with predicted gnomAD-pLOF variants that overlap predicted and experimental Z-flipons and that are also present in the Morbid OMIM gene set.
